# The cost of efficiency in flexible neural representations

**DOI:** 10.64898/2026.05.18.722885

**Authors:** Wenhao Dang, Peiyu Chen, Christos Constantinidis

## Abstract

Working memory depends on the flexible representation of stimulus information in neural activity, which changes dynamically depending on task. Stimulus transformations are thought to be efficient in use of neural resources and optimal for task performance. However, these transformations are often opaque, and efficiency may conflict with optimal performance. Here we show that in a working memory task requiring selective recall of one of two stimuli based on a context cue, the prefrontal cortex of two male monkeys prioritized efficiency by overwriting information within a shared neural subspace rather than maintaining distinct subspaces for each stimulus. In neural activity and recurrent neural networks such efficiency incurs a cost, in that efficient representations are more prone to errors. Conversely, stimulation of the cholinergic forebrain which improves behavior altered this default mechanism by encoding distinct contexts in higher dimensions. These findings demonstrate a fundamental tradeoff between efficiency and effectiveness in flexibly updating working memory.

## INTRODUCTION

The representation of stimuli in the activity of neuronal populations can change dynamically during the course of a trial, depending on what information is critical for the execution of a task^1, 2^. A variety of stimulus manifold transformations have been described during performance of various cognitive tasks, such as projections to different subspaces or rotations^3, 4, 5^. Orthogonal rotation of a stimulus representation, in particular, has been proposed as a potential mechanism for reducing interference between multiple sensory and memory representations, protecting the memory of an initial stimulus from the interference of a subsequent stimulus presentation, or stimuli that may be represented in different subspaces when used in different task contexts^6, 7^. In multi-item sequence tasks, items are encoded in distinct subspaces based on their presentation order and subsequently reorganized in a context-dependent manner^8, 9, 10^.

It is commonly assumed that stimulus transformations during the performance of cognitive tasks are efficient in terms of the use of neural resources and optimal for performing the task at hand, thus achieving effectiveness^5, 11, 12^. Efficient coding has been traditionally used to describe sensory stimulus representations in neural activity^13, 14, 15^. More recently, the principle has been extended to optimal implementations of cognitive computations^16^. However, it has been less appreciated that efficiency and effectiveness can be at odds. This is a well understood dichotomy in business management, which identifies efficiency in terms of minimizing use of resources and effectiveness in ensuring achievement of the desired objective ^17^, though these concepts are often confused in neuroscience^12^.

We were thus motivated to investigate the representational geometry of population activity in the prefrontal and posterior parietal cortex when subjects are required to remember one of several stimuli under different contexts. In such a task, multiple potential strategies, which differ in their efficiency and effectiveness, are viable at the algorithmic level. We examined what strategies were followed by neural activity. We also relied on artificial recurrent neural networks (RNNs) under different training regimes to evaluate the efficiency and effectiveness of these transformation. Finally, we performed perturbation of this activity with deep brain stimulation to draw causal conclusions on how task performance relates to the efficiency of stimulus representations.

## RESULTS

Two male monkeys (*Macaca mulatta*) were trained to perform the Remember 1^st^ (R1) – Remember 2^nd^ (R2) task (Fig. 1 A). The monkeys viewed two stimuli presented in sequence, with a delay period intervening between them. After another delay period, the fixation point turned off, and they had to make an eye movement in either the first stimulus, or the second, depending on the color of the fixation point (white or blue). Monkeys were generally able to choose the correct stimulus, however smaller target-distractor distances were associated with lower accuracy (Fig. 1B-C) and a systematic attraction of the reported location toward the distractor was observed (Fig. 1D). A total of 433 neurons were recorded (Fig. S1) from areas 8 and 46 of the dorsolateral prefrontal cortex (PFC, N = 198, with 12,041 trials for R1 and 11,520 trials for R2) and areas 7a and LIP of the posterior parietal cortex (PPC, N = 235, with 14,979 trials for R1 and 13,989 trials for R2).

**Figure 1.**
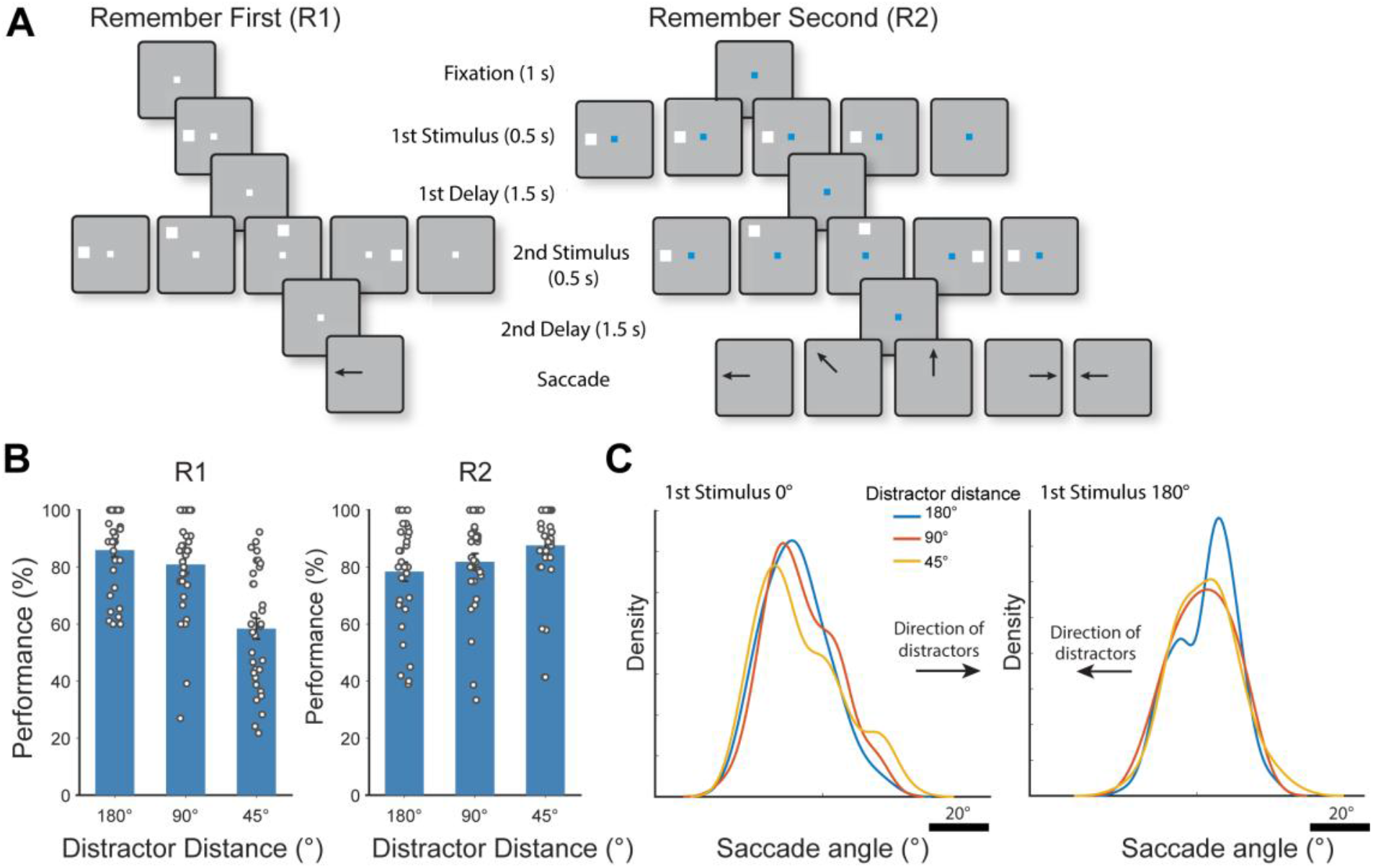
Neural recording areas and the context-dependent working memory task paradigm. (A) Diagram of the Remember 1^st^ -Remember 2^nd^ tasks. Frames illustrate the sequence of events in the Remember 1^st^ (left) and Remember 2^nd^ (right) contexts. (B) Behavioral performance of the monkeys by context and target-distractor distance. (C) Saccadic endpoint distribution in Remember 1^st^ context for different stimulus conditions, sorted by first stimulus location and distractor distance The monkeys’ saccade report locations were systematically biased by the distractor.

### Three potential models of context-dependent working memory

At the level of neural activity, multiple strategies could in principle support successful task performance. For example, the brain might retain only the context-relevant stimulus. Alternatively, it could maintain separate representations of both stimuli and retrieve the context-relevant information at the time of report. In the former case, neural activity might reuse a single subspace, flexibly storing stimuli according to the context cue. In the latter case, both locations might be stored in distinct subspaces, organized by either their temporal order in the trial or their task relevance based on the context cue. We therefore distinguish between three models of neural representations to encode the first and second stimulus locations in the Remember 1^st^ – Remember 2^nd^ task, representing discrete cases along a continuum of possibilities that differ in terms of their efficiency, as outlined below and in Fig. 2A:

**Figure 2.**
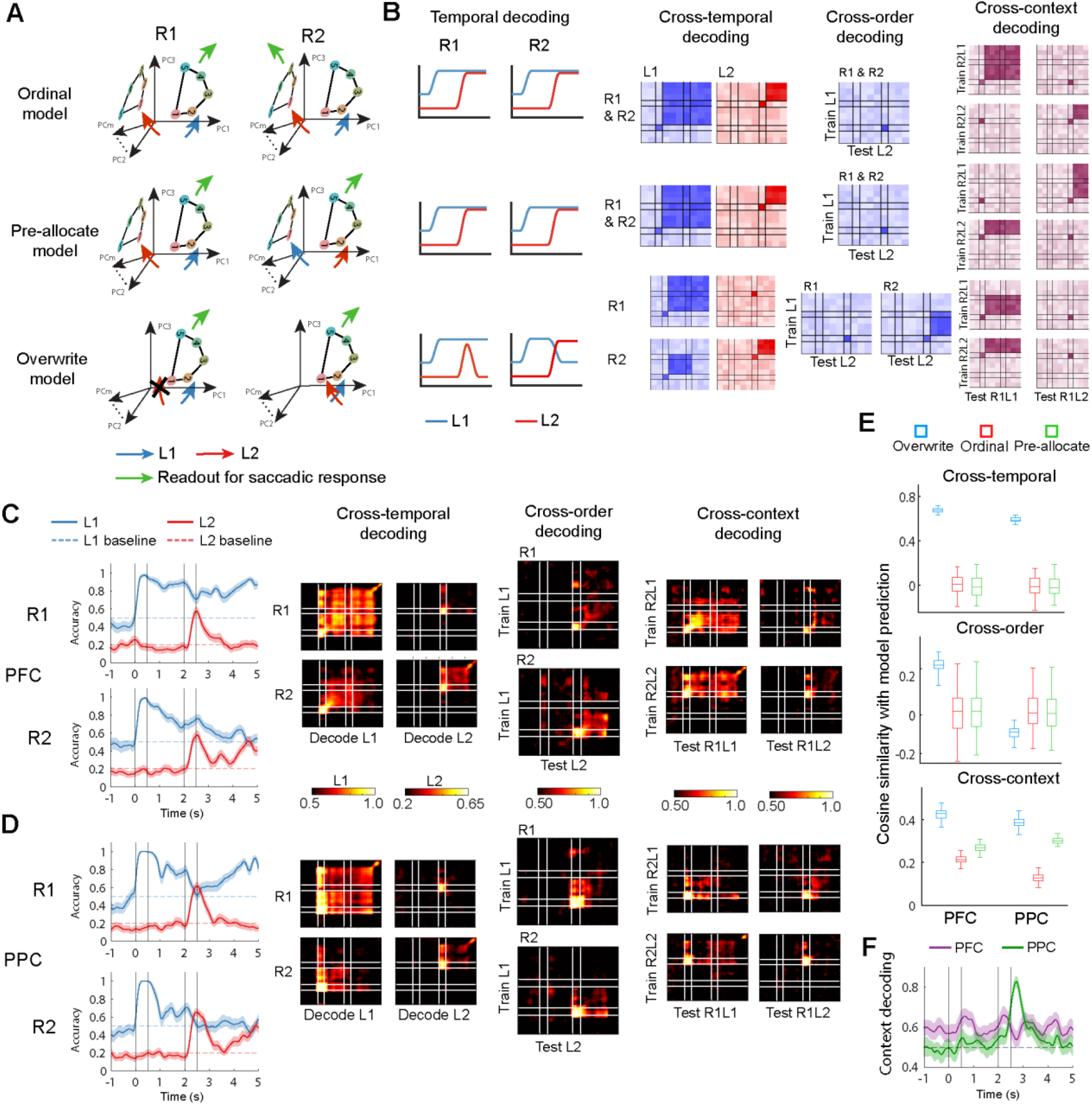
Monkeys utilize context-dependent working memory by gating and overwriting representations within a shared subspace. (A) Schematic illustration of three models for performing a context-dependent working memory task in low-dimensional subspaces. Representation of five stimulus locations (colored dots) used in the context of the Remember 1st and Remember 2nd task are shown in different PCA subspaces. The Pre-allocate model (middle left) suggests that representations are encoded in subspaces defined by task relevance, with task-relevant information consistently occupying the same subspace across different contexts. The Overwrite model (bottom left) proposes that stimulus representations may be either stored or overwritten in the same subspace, contingent on context. Diagrams on the right of each model illustrate the predicted decoding performance for each scenario. (B) Predicted decoding performance across time, across specific stimuli, and across Remember 1^st^ versus Remember 2^nd^ contexts for the three models. Each matrix element represents the decoding accuracy of a classifier trained at one time point and tested at another time point. (C, D) Empirical decoding performance of the first and second stimuli in (C) the prefrontal cortex and (D) the posterior parietal cortex. The first stimulus had two possible locations (chance = 0.5), whereas the second had five (chance = 0.2). For the cross-order and cross-context decoding analyses, we used the two shared locations across the two stimuli, yielding a chance level of 0.5. Cross-order decoding and cross-context decoding show that the information can be shared across order and task in the PFC. Solid lines show the empirical decoding accuracy, while dashed lines indicate chance level. Vertical lines separate a trial into five time epochs, as indicated in Figure 1: fixation, first stimulus, first delay, second stimulus, and second delay periods. (E) Cosine similarity between empirical decoding results and the predictions of the three models. The decoding profiles of the Overwrite models have higher cosine similarity with the PFC empirical data. The Overwrite model is also more similar to the PPC data except for cross-order decoding. (F) Context decoding performance (i.e., distinguishing Remember 1^st^ vs. Remember 2^nd^ contexts) in the prefrontal cortex and the posterior parietal cortex. The time point of zero refers to the onset of the first stimulus. Abbreviations R1: Remember 1^st^; R2: Remember 2^nd^; L1: the first stimulus location; L2: the second stimulus location.

1. *Ordinal model*: This model posits that both locations are represented in subspaces defined by their temporal order; such representations have been shown to emerge in tasks that require sequential working memory, i.e. remembering multiple stimuli and their sequence of presentation^18^. For each context, the targeted location is read out from either the first- or the second-order subspace.
2. *Pre-allocate model*: If it is known that only one stimulus needs to be remembered, an alternative model is possible, that assumes that representations are stored in context-specific subspaces, one for targets and one for distractors (non-remembered stimuli)^1, 19^. In this framework, the first stimulus in the Remember 1^st^ task and the second stimulus in the Remember 2^nd^ task are stored in the target subspace, while the other stimulus is stored in the distractor subspace.
3. *Overwrite model*: In both the Remember 1^st^ and Remember 2^nd^ contexts, the first location is initially stored in this shared subspace. The neural dynamics diverge after the presentation of the second location: in the Remember 1^st^ context, the second location does not enter the shared subspace, preserving the original memory trace; in the Remember 2^nd^ context, the second location overwrites the memory trace of the first location upon its presentation.

Intuitively, the Overwrite model is the most efficient among the three models under comparison, since the Ordinal and Pre-allocate models require an additional subspace, which consumes more resources for maintenance and manipulation. However, it is also the least effective from the perspective of potential interference of information in the same coding space. More formally, efficiency in cognitive computations has recently been defined as an optimization problem to minimize the sum of neural activity and dendritic weight^16^:

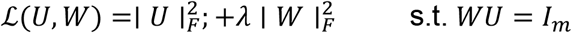

Where *m* is the dimension of the variable to reconstruct and *λ* > 0 is a tradeoff hyperparameter determining the relative contribution of neural activity and weight to energy cost. It can be proven (see Supplementary Information for full details) that:

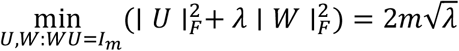

In other words, the minimum cost scales linearly with the dimensionality of the task variables maintained in memory and a model that maintains fewer variables (such as the Overwrite) will be more efficient than those that maintain more (Ordinal and Pre-allocate).

In the same framework, we can also evaluate the effectiveness of the different models, defined as the inverse of the Mean Squared Error (MSE) of their decoding output. Consider a representation *r* ∈ ℝ^*N*^ that must allow linear decoding (with decoding matrix *U* ∈ ℝ^*N*×*m*^) of some task variable *z* ∈ ℝ^*m*^ with zero error. Focusing on the critical case of the Remember-2^nd^ context, where overwrite must erase the first memory, the end-of-trial representation is

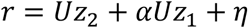

Where *α* is residual leakage of the first stimulus after overwrite (*α* = 0 is perfect overwrite; *α* ≠ 0 is incomplete erasure / imperfect gating), and 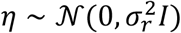 is activity noise. With a decoder *W* satisfying *WU* = *I*, the decoded output is

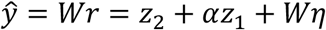

The MSE (inverse of effectiveness) is given by

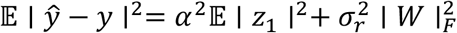

We can then compare effectiveness to models that store both subspaces (Ordinal and Pre-allocate) models. In the Pre-allocate model, with ideal subspace separation the same equations are expressed as,

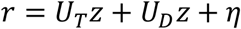

By definition, *W*_*T*_*U*_*D*_ is zero, so there is no *α*^2^𝔼 ∣ *z*_1_ ∣^2^ interference term in this case. That means that with any nonzero overwrite leakage, the Overwrite model will be less effective i.e. less precise in decoding the stimulus.

### The efficient Overwrite model best predicts neural coding of context-dependent working memory

To test which model best describes neural activity, we examined model predictions for decoding accuracy (in this section), and for representational geometry (in the following section), and compared these predictions with the empirical data. The three models make distinct predictions for temporal decoding accuracy, as well as cross-temporal, cross-order, and cross-context decoding performance both across and within contexts (Fig. 2B). In terms of temporal decoding, both the Ordinal and Pre-allocate models preserve information of both stimuli, maintaining high decoding accuracy after their presentation, regardless of the contextual cue. Thus, both models yield the same decoding performance for both locations across contexts. On the other hand, in the Overwrite model, the first location is encoded invariantly in both contexts before the second stimulus appears. However, the introduction of the second stimulus has different influences on the existing memory representations between contexts. Specifically, in the Remember 1^st^ context, the representation of the second location diminishes after the second stimulus appears. In the Remember 2^nd^ context, the second stimulus overwrites the representation of the first location, leading to a decrease in decoding accuracy.

In cross-temporal decoding analysis, a decoder of stimulus location is trained at one time point and tested at all time points under the same context (e.g. the first stimulus of the Remember 1^st^ condition). This analysis tests how persistent the information is for two stimuli used in the task. The Ordinal and Pre-allocate models predict that the location could be decoded across time after the stimulus shown up, as the representations of the two stimuli remain in their own separable subspace throughout their delay period. In contrast, the Overwrite model predicts that the cross-time decoding generalization for both spatial locations are context-dependent. Specifically, the code used for the first stimulus should persist throughout the trials under Remember 1^st^ context but diminish after the second stimulus presentation under the Remember 2^nd^ context. Conversely, the decoding performance for the second stimulus should drop sharply when the second stimulus is no longer on the screen for the Remember 1^st^ context.

In cross-order decoding analysis, a decoder of stimulus location is trained on one time point and tested on the location of the other stimulus used in the same trial for all time points in the same context (e.g. the decoder is trained with the first stimulus in the Remember 1^st^ context, and tested on the location of the second stimulus in the Remember 1^st^ context). This analysis tests the similarity for neural representation for the first vs. the second stimulus in the same context. Neither the Ordinal or Pre-allocate models can support generalization in cross-order decoding analysis since the locations of the first and second stimuli are stored in two different subspaces. In contrast, the Overwrite model exhibits high cross-order decoding performance, because the second stimulus overwrites the first in the shared subspace, and representations across stimulus orders become generalizable.

The three models also predict distinct cross-context generalized patterns. The Pre-allocate model maintains the same decoding pattern for each location since both stimuli are stored in the same subspace across contexts. In the ordinal model, the target location (first location of the Remember 1^st^ and second location of the Remember 2^nd^) can be generalized across their corresponding delay period, so as two distractors. For the Overwrite model, the first location during the first delay period for both tasks and the second location during the second delay period could be generalized owing to the dynamic overwriting mechanism.

To evaluate which model best captures the neural coding dynamics, we performed cross-temporal, cross-order, and cross-context decoding to neural population activities from the PFC and the PPC of monkeys performing the Remember 1^s^-Remember 2^nd^ task (Fig. 2C-D). In cross-temporal analysis, most consistent with the Overwrite model, we observed context-dependent modulation of location information for both stimuli. Specifically, in both PFC and PPC, the average decodability for the first stimulus dropped significantly in the second delay for subjects under the Remember 2^nd^ context compared to the Remember 1^st^ context (PFC from 77.5% to 59.5%, *t* (18) = 17.0, *p* = 1.5×10^-12^, two sample t-test; PPC from 70.2% to 51.7% in the Remember 1^st^ context, *t* (18) = 20.7, *p* = 5.5×10^-14^, two sample t-test), while the decodability for the second stimulus was significantly higher in the second delay under the Remember 2^nd^ context compared to the Remember 1^st^ context (PFC from 20.9% to 30.4%, *t* (18) = 6.0, *p* = 1.1×10^-5^, two sample t-test; PPC from 22.3% to 28.7% in the Remember 1^st^ context, *t* (18) = 6.8, *p* = 2.3 ×10^-6^, two sample t-test). Furthermore, we hypothesized that if the “overwrite” strategy were used, behavioral errors would be accompanied by reduced context-dependent gating of information. Consistent with this prediction, such a pattern was observed in the PFC, especially in the *Remember 1st* context, where most error trials occurred (Fig. S6). Despite a similar pattern in temporal decoding in PPC, some differences between areas were also present. More stimulus location information could be decoded during both delay periods from the PFC than PPC, and even though there were fewer PFC neurons in the current dataset. Under the Remember 1^st^ context, the average decoding accuracy for the first stimulus across the two delays was 81.6% in PFC and 73.5% in PPC. In the Remember 2^nd^ context, the decoding accuracy for the second location in the second delay was 30.4% in the PFC and 28.7% in PPC. Moreover, we found that decodable information was more susceptible by the second stimulus in the PPC, leading to more dynamic decoding results. In the Remember 1^st^ context, the decoding performance in PPC sharply dropped from 78.0% in the first delay to 63.0% right after the presentation of the second stimulus, while in PFC it only dropped from 85.9% to 78.6%. Interestingly, under this context, the decoding performance in PPC bounced back to 80.2% toward the end of the second delay, comparable to the performance in the first delay. In contrast to the more stable representation in PFC, this result suggests that the mechanism of sustaining working memory in PPC is different from the PFC, or that information about the stimulus was fed into the PPC from other areas after distraction.

We similarly tested the agreement of neural data with the three model predictions in terms of stimulus-encoding subspaces across orders (i.e. first vs second stimulus), as well as across contexts (i.e. Remember 1^st^ vs Remember 2^nd^, Individual subject results see Fig. S4). In PFC, cross-order decoding shows that the code used for the first location in the first delay epoch could be generalized for the second location in the second delay when the second location is to be remembered (Fig. 2C, middle heatmap). Similarly, cross-context decoding revealed that the code used for the second stimulus location under the Remember 2^nd^ context, could decode the first location in the Remember 1^st^ context. Together, these results suggest there exists one common working-memory subspace used across all delay epochs to maintain the relevant location information in this task.

To quantify how well the neural data fit each model, we computed the cosine similarity between the models’ decoding matrices and neural decoding matrices across all decoding analyses (Fig. 2E). The Overwrite model showed the highest correlation with neural data across all predictions in PFC. For PPC, although the Overwrite model was the best predictor for cross-temporal decoding and cross-context decoding across the three models we compared, it failed to accurately depict cross-order decoding results. Though the Overwrite model better predicts the decoding performance of the PFC and PPC, the agreement with the model was not absolute and could best be characterized as “partial overwriting” of stimulus coding space.

Furthermore, the PPC and PFC behaved qualitatively differently in maintaining context information. Task context information could be stably decoded from the PFC throughout the trial, though the performance was relatively low (Fig. 2F). In PPC, there was a transient peak in context information during the early second delay epoch, but the decoding performance was not better than chance during other periods of the trial. Such information in the PPC could serve to gate access to the shared common space found in the PFC.

### Representational rotation and coding reorganization also support the efficient Overwrite model

Besides examining whether stimuli occupy a shared or distinct subspace across task epochs, we can reveal how neural representations distinguish the two contexts by determining the rotation of stimulus locations between the Remember 1st and the Remember 2nd context for the same epoch. We calculated the rotation angles between the subspace defined by the stimuli when presented in the first stimulus interval and the subspace defined by the stimuli in the second stimulus interval. We repeated this analysis for the delay intervals, and separately for the Remember 1st and Remember 2nd contexts. The three theoretical models make distinct predictions about these rotation angles between contexts (Fig. 3A and Fig. S2) and we exploited them to independently test the similarity of the models with neural data (Fig. 3A). To do this, we calculated the subspace rotation angles between the Remember 1^st^ and Remember 2^nd^ contexts, for each epoch. Specifically, we applied principal component analysis (PCA) on the neural response matrix for each epoch, and defined rotation angle as the principal angle between the subspaces containing the representation for each context (Remember 1^st^ vs Remember 2^nd^; see Methods). In our study, the first three principal components (PCs) explained more than 72% of the variance in the response matrix across all analyzed periods and locations, indicating that neural activity could be effectively represented within a low-dimensional subspace (Fig. S3).

**Figure 3.**
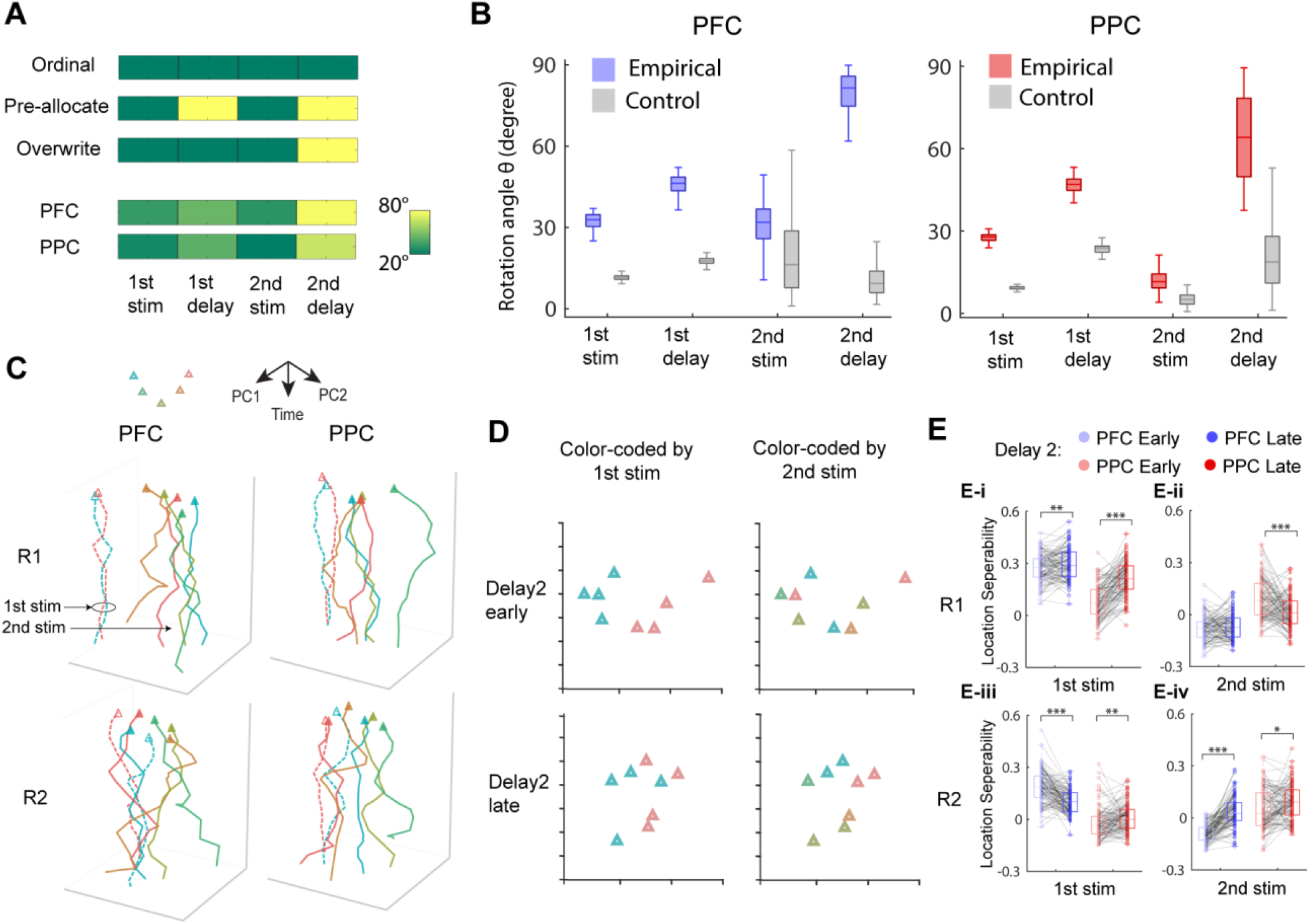
Context-dependent working memory drives subspace rotation and representation reorganization in PFC and PPC. (A) Predicted and empirical rotation angles in four trial epochs for the three models—ordinal, pre-allocate, and overwrite—as well as the prefrontal cortex and the posterior parietal cortex. (B) Subspace rotation angles between the Remember 1^st^ and the Remember 2^nd^ contexts across task epochs. Rotation angle variability was assessed via bootstrap resampling of trials and was then compared to a control condition generated from a synthetic dataset (see Methods). (C) Beta-space representation of the first (dashed lines) and second locations (solid lines) during the second delay in PFC (left) and PPC (right). (D) Visualization of spatial code reorganization of the PFC. Neural responses during early (250 ms post-second stimulus offset) and late (1250 ms post-second stimulus offset) phases of the second delay epoch in the Remember 2^nd^ context were projected into low-dimensional space. Responses were color-coded based on the spatial location of either the first or second stimulus, using the same color-location correspondence shown in panel C. During the second delay period, separability of the first stimulus representation decreased, while separability of the second stimulus representation increased. (E) Quantification of geometrical separability for the first and second stimuli during the second delay epoch in Remember 1^st^ and Remember 2^nd^ contexts for both the prefrontal cortex and the posterior parietal cortex. *, **, *** indicate *p* < 0.05, *p* < 0.001, *p* < 0.0001, evaluated using a two-tailed paired t-test.

Neural subspace rotations between the Remember 1^st^ and Remember 2^nd^ contexts followed a similar pattern for the PFC and PPC across different trial epochs. Relatively small rotation angles were found during the two stimulus presentations, but still significantly greater than rotation angles expected by chance (*p* < 0.001, permutation test). Stimulus rotation angle increased during the first delay period, and near orthogonal rotations were found in the second delay (Fig. 3B). Alternative methods for measuring subspace alignment based on dimension reduction on lasso regression coefficients, Canonical Correlation Analysis (CCA) and variance accounted for (VAF) ratio produced similar results (see Fig. S7 and Methods for implementation details). Small rotation angles up until the second delay period best aligned with the predictions of the Overwrite model (Fig. 3A), as neuron populations perform identical operations across contexts up to that point. The high rotation angle in the second delay period is due to the critical context-dependent operation in the Overwrite model performed at that interval (gating or overwrite, for R1 and R2, respectively, as hypothesized).

To further explore the working memory manipulation during the second delay, which distinguished two contexts and resulted in high subspace rotation angles, we visualized the neural response in a low dimensional subspace across the time course of the trial. Specifically, we calculated the firing rates of each neuron in a time-resolved fashion and then applied Lasso-regularized regression to extract components corresponding to the first and second stimuli, respectively, from the neural activity. The fitted beta coefficients at each time point formed the “beta-space” representations for the two locations. The beta-space representational trajectories for the first and second locations were visualized using principal component analysis (PCA), with the first two principal components shown in Fig. 3C. We hypothesized that, in the Remember 2^nd^ context, if the two stimuli occupy a common coding space, the neural trajectories for R1 and R2 would be intermingled. By contrast, in the Remember 1^st^ context, the trajectories should form two distinct clusters, because one of the stimuli is not persistently represented.

These beta-space representations indeed revealed distinct trajectories patterns for the Remember 1^st^ and the Remember 2^nd^ contexts. In the Remember 1^st^ context, the trajectory for the second location remained distant from that of the first location. In contrast, in the Remember 2^nd^ context, the two trajectories appeared to overlap or intersect, suggesting that the second location occupied a neural space that had previously represented the first location. This pattern confirms that in the Remember 2^nd^ task, the second location becomes the new target, replacing the first one. Conversely, in the Remember 1^st^ task, the second location is represented as a distractor, remaining well separated from the target representation. These results align well with our decoding finding that in both contexts, stimulus-location codes reside in the same space with a dynamic overwriting and gating mechanism.

We further hypothesized that in the Remember 2^nd^ context, we should be able to observe the neural code for the second stimulus gradually replace the code for the first stimulus. To directly test this hypothesis, we visualized the neural response of the early (250 ms post-second stimulus offset) and late (1250 ms post-second stimulus offset) second delay epoch under the Remember 2^nd^ context in low dimensions, shown color coded with reference to either the first or second stimulus in Fig. 3D. As we predicted, for the critical, Remember 2^nd^ context (blue points in Fig. 3E-iv), the PFC neural representation optimized the separation of the first stimulus early in the second delay period, but this separation decreased significantly towards the end of the delay period (*t* (99) = 7.0, *p* = 3.4×10^-10^, two-sided paired t-test). In contrast, the separation of the second stimulus was low early and increased significantly in the latter part of the delay period (*t* (99) = 8.7, *p* = 7.6×10^-14^, two-sided paired t-test). No such pattern of reorganization was present (or expected), in the Remember 1^st^ context (Fig. 3E-i,ii). The separation of the first stimulus in the Remember 2^nd^ task for the PPC did not appear to diminish and rather increased (*p* < 0.001, permutation test), even though this stimulus was no longer relevant (red points in Fig. 3E-iv). These results point out an efficient representation of stimulus information, specifically for the PFC.

In summary, besides the efficient working memory maintenance, the subspace rotation analysis points to a low-cost working memory manipulation strategy. For most of the trial (up to the second delay), the neural population maintains only a modest rotation between the two contexts, implying that neuron activity undergoes slight changes between contexts. The critical context-dependent manipulation only occurs in the second delay, evidenced by the high subspace rotation angle and different coding reorganization between two contexts. This pattern suggests that the network conserves resources by performing a common computation for as long as possible and engaging context-dependent transformations only when strictly necessary.

### RNNs under rich training regime gravitate to efficient Overwrite geometry

The empirical results point out to some transformations of neural representations at different task epochs and contexts. We wondered what factors could determine how a neural network solves a context-dependent working memory problem when potentially multiple solutions exist. We therefore constructed artificial neural networks that allow the generation of diverse representation geometries for the same task ^20^. Specifically, we trained recurrent neural networks (RNNs) with gated recurrent units (GRUs) under different learning regimes^21^ to perform the same Remember 1^st^ – Remember 2^nd^ working memory task.

Previous research has demonstrated that changing the variance of weight initialization leads to qualitatively different internal representations for the same task ^20, 22, 23^. In particular, the “rich training” regime, characterized by small initial weight variance, yields low-dimensional and structured representations. In contrast, the “lazy training” regime, characterized by large initial variance, randomly projects inputs into higher-dimensional spaces that support more separable representations. To systematically evaluate the representations under different training regimes, we trained RNNs with weight initialization ranging from 0.2 to 3.0 in steps of 0.4 (Fig. 4A; see methods for training details). We define the rich regime as weight initialization ≤ 1.0, and the lazy regime as > 1.0. As expected, richly trained RNNs exhibited lower dimensionality than lazily trained ones, quantified by the participation ratio (Fig. S5A).

**Figure 4.**
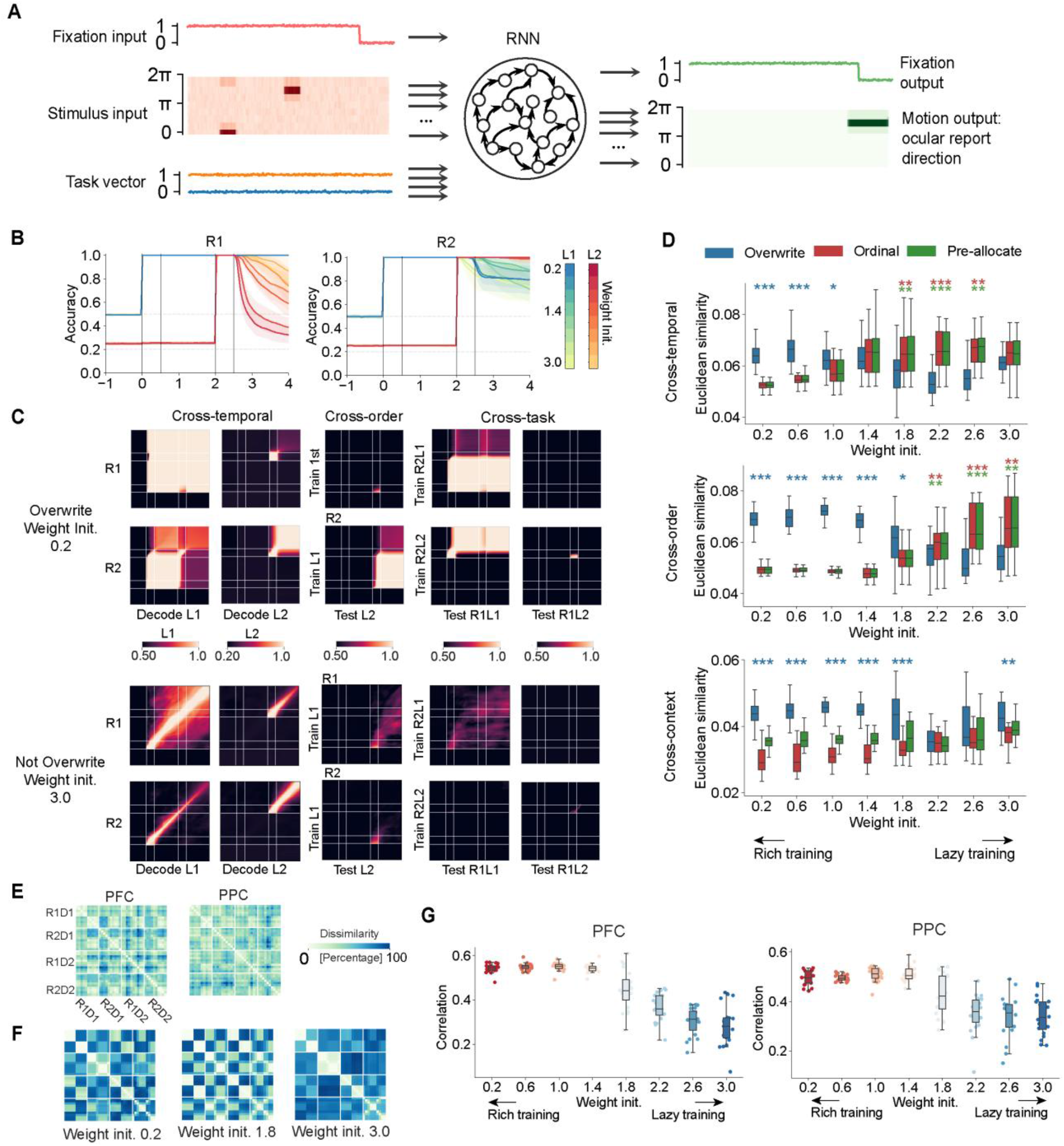
Richly trained RNNs replicating Overwrite representational geometry. (A) Diagram of the inputs and outputs of the recurrent neural networks (RNNs). (B) Temporal decoding of RNNs under rich (small initial weight values – dark hues) and lazy training regimes (large initial weight values – light hues). (C) Two example decoding profiles of cross-temporal, cross-order, and cross-context decoding of RNNs representations under rich (upper) and lazy (lower) training regimes. (D) Euclidean similarity between empirical decoding results and the predictions of the Ordinal, Pre-allocate, and Overwrite models. (E, F) Representation dissimilarity matrices are calculated for neural data (E), and the RNNs (F). (G) The Pearson correlation *r* between representation dissimilarity matrices of empirical neural data and the RNNs. *: *p* < 0.05, **: *p* < 0.01, ***: *p* < 0.001, evaluated using a two-sided paired t-test to compare whether the best-fitting model exhibited significantly higher similarity than the second-best model. Colors indicate the best-predicting model (blue: Overwrite; red: Ordinal; green: Pre-allocate). Note that the Ordinal and Pre-allocate models share identical decoding profiles (Fig. 2B) for cross-temporal and cross-order decoding, and thus, both red and green colors are being used here. Abbreviations R1: Remember 1^st^; R2: Remember 2^nd^.

After training, we found that the decoding accuracy of the distractor (non-remembered) stimulus location decreased during the second delay period (Fig. 4B), as predicted by the Overwrite model. The richly trained RNNs exhibited lower decoding accuracy of the distractor location during the second delay for both the Remember 1^st^ (rich < lazy, *t* (158) *=* -63.37, *p* = 0.0, two-sided t-test), and the Remember 2^nd^ context (rich < lazy, *t* (158) = -4.73, *p* = 2.3 × 10^-6^, two-sided t-test). We further performed cross-temporal, cross-order, and cross-context decoding (Fig. 4C), and calculated their Euclidean similarity (Fig. 4D) to the theoretical models described in the previous sections (see Fig. 2). Only RNNs trained under the rich training regime significantly matched the overwrite model across all three decoding profiles (Overwrite > Others, *p* < 0.05 for all decoding profiles, two-sided paired t-test; Fig. 4C, top panel for weight initialization = 0.2). In contrast, when the weight initialization was set to 3.0, the resulting representations more closely resembled a dynamic coding model, where each time point has its own unique codes (Fig. 4C, bottom panel).

Furthermore, we examined whether the representational geometry of richly trained RNNs, which exhibited “Overwrite”-like representations, more closely resembled empirical neural population activity. We constructed task-based representations for all task conditions and two delay periods using the representation dissimilarity matrix (RDM)^24^, as shown in Fig. 4E for neural data and Fig. 4F and Fig. S8F for RNNs. This result aligns with the “Overwrite” model where the shared subspace ensures one low-dimensional manifold is enough for the task, while the other two models require two. To quantify the similarity between RNNs and empirical neural RDMs, we computed the correlation between the two (Fig. 4G). We found that the average correlation declined as the weight initialization increased beyond 1.4. Richly trained RNN showed significantly higher correlation with neural data from PFC (*t* (158) = 13.21, *p* = 3.3 × 10^-24^, two-sample t-test) and PPC (*t* (158) = 10.26, *p* = 4.0 × 10^-18^, two-sample t-test). These findings offer evidence that the Overwrite model represents an efficient coding strategy for context-dependent working memory in low-dimensional spaces, consistent with the empirical neural representation.

### Subspace rotation increased with performance in richly trained RNNs

The RNN models allowed us to investigate the implications of one more empirical finding: Subspace rotations between contexts intuitively appear to offer an advantage in reducing the interference between stimuli from different contexts (Fig. 3B). We therefore tested the hypothesis that rotations between contexts allowed better performance. The representations of RNNs can also be embedded within low-dimensional subspace, with the first three PCs capturing more than 77% of the variance (Fig. S8B). Under the rich training regime, rotation angles were small initially and increased significantly across all task phases after training (Fig. 5A & S8D; *p* < 0.001, two-sided paired t-test across all task epochs). After training, the relative differences in rotation angles across phases resembled the empirical findings, with the largest rotation angles observed during the second delay period and lower, but non-zero rotation angles present during other periods. By contrast, lazily trained models exhibited high initial rotation angles, but they did not change significantly with training (Fig. S8D). These results imply that subspace rotation plays a meaningful role in supporting context-dependent working memory.

**Figure 5.**
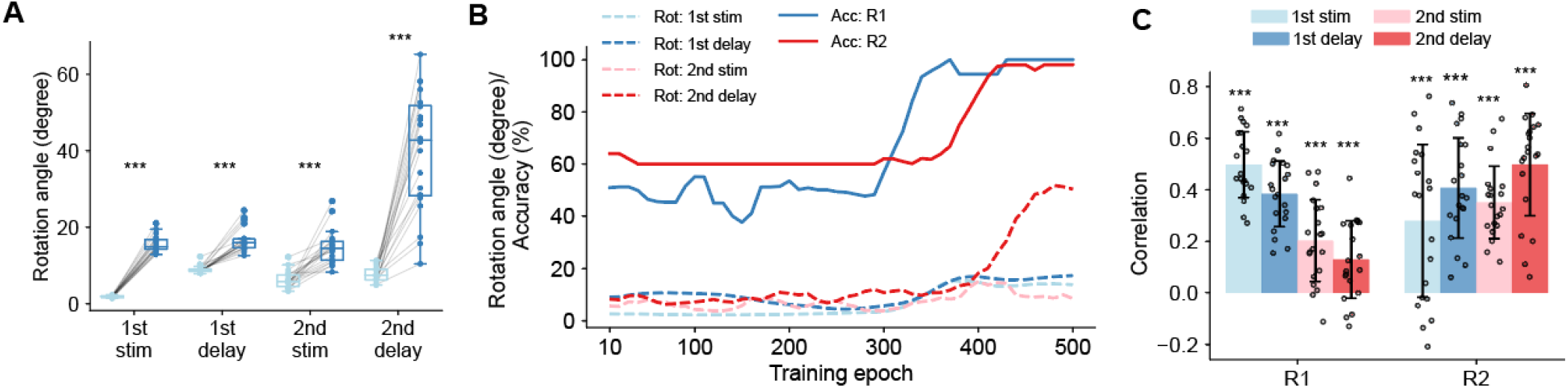
Subspace rotation increased with the performance at richly trained RNNs. (A) Beta-space subspace rotation angles of RNNs’ representations before and after training. (B) An example of increased subspace rotation and task performance over the course of training. (C) Pearson correlation (*r*) between the subspace rotation angles and training performance (accuracy) in Remember 1^st^ (R1) and Remember 2^nd^ (R2) contexts. ***: *p* < 0.001, evaluated using a two-sided paired t-test for (A) and two-sided t-test for (C).

To examine whether changes in subspace rotation are linked to task learning, we computed Pearson correlations (*r*) between task performance and subspace rotation angles (Fig. 5B-C & S8D). To control for training duration, we regressed out the number of training epochs from both rotation angles and performance scores, as both metrics tend to increase over the course of training. For rich training, performance on both Remember 1^st^ and Remember 2^nd^ contexts was significantly correlated with subspace rotation angles for all phases (all *p* < 0.001 using two-sided t-test to evaluate *r* > 0). This relationship was not significantly observed under lazy training (Fig. S8D).

The RNN analysis provided two additional, non-trivial, insights. First, performance in the Remember 1^st^ task was most strongly associated with subspace rotation during the first stimulus and first delay periods, while Remember 2^nd^ performance correlated more with rotation during the first and second delay periods. Secondly, many prior studies have emphasized orthogonality between stimulus representations as a means of reducing interference^6, 7, 11^. The results of Fig. 5A suggest that even more modest rotations provide performance advantages. In our data, we found relatively low, but significantly higher than chance rotations in the first 3 epochs (Fig. 3B,1^st^ stim, 1^st^ delay, 2^nd^ stim). The RNN results suggest that the simultaneous maintenance of stimulus and context information (which we saw in neural data, in Fig. 2F) may contribute to this modest rotation and potentially could play an important role in task learning and execution.

### Overwrite RNNs are efficient but less effective than Ordinal and Pre-allocate RNNs

To test directly whether different representational geometry imposes a tradeoff between efficiency and effectiveness, we constructed Ordinal and Pre-allocate versions of the RNN models that explicitly maintained separate subspaces for different locations, as Fig 2A. In both cases, the recurrent architecture was identical to that of the Overwrite model, but the readout structure was modified to add two Heads, A and B, together with an orthogonality penalty that led the two heads to span distinct subspaces. In the Ordinal model, Head A and Head B were trained to decode the first and second stimuli, respectively. In the Pre-allocate model, Head A and Head B were trained to decode target and distractor information, respectively (Fig. S5A). By contrast, the Overwrite model, described in the preceding section, used a single readout head to decode the task-relevant representation (Fig. 4A).

We first confirmed that our RNNs captured the conceptual models we proposed. In particular, they faithfully reproduced the predicted decoding patterns (Fig. 2; Fig. S5B), and the “overwrite” RNN additionally recapitulated the empirical behavioral results observed under noise (Fig. S5E). We then compared the three network classes using various measures of efficiency and effectiveness. In the case of efficiency, energy and effective dimensionality were calculated. Both the Ordinal and Pre-allocate networks showed significantly higher activity energy and higher effective dimensionality than the Overwrite networks (Fig. S5C), indicating that maintaining two separable subspaces requires greater representational resources, consistent with our theoretical hypothesis. However, this additional cost was associated with more effectiveness, which was evaluated by the robustness to noise. When Gaussian noise was added to the stimulus input during evaluation, both the Ordinal and Pre-allocate networks achieved higher task accuracy than the Overwrite networks in both the Remember 1^st^ and Remember 2^nd^ contexts, with the difference becoming more pronounced as noise increased (Fig. S5D-E). Thus, models that segregate information across distinct subspaces are less efficient, because they rely on greater activity and higher-dimensional representations, but they are more effective in preserving performance under noise perturbation. Conversely, the Overwrite solution provides a more efficient representational strategy, but this efficiency makes it more vulnerable to noisy interference.

### Errors exhibit lower PFC rotations but no less adherence to the Overwrite model

Analysis so far showed that PFC, and to a lesser extent PPC, utilized an efficient, low dimensional method for encoding information. We also revealed that in our RNN model alternative coding schemes exist as long as the network operates in a different dynamic regime. To test their significance, we examined error trials. First, we examined the adherence of neuronal activity to the Overwrite, Ordinal, and Pre-allocate models (Fig. 6A). Error trials were fewer than correct and absent in some conditions (overall performance was 82.5%±1.4% and 88.6%±1.3% for the two subjects, respectively). We therefore compared neural activity from correct trials alone with activity from both correct and error trials. For both correct and error trials, the Overwrite model provided the best fit; adding error trials in the analysis did not change the order of model-data similarity, for either the PFC or PPC. The transformation achieved by the Overwrite model suggests efficiency from the perspective of neuronal representations. For error trials, the Overwrite model best described the prefrontal neural activity data.

**Figure 6.**
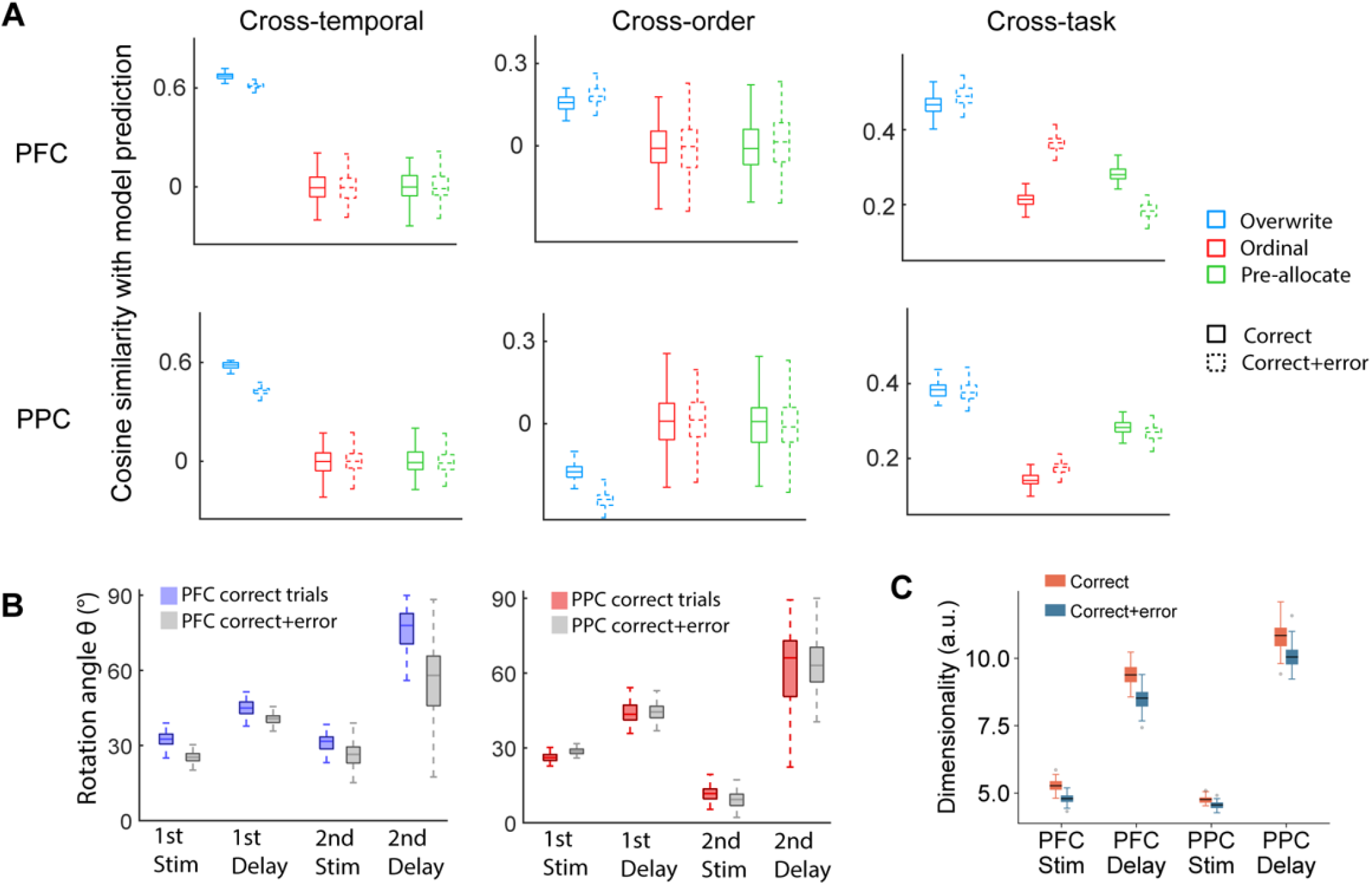
Error trials. (A). Cosine similarity between empirical decoding results and the predictions of the ordinal, pre-allocate, and overwrite models. (B) Subspace rotation angles between R1 and R2 contexts in each epoch for PFC (left) and PPC (right), using correct trials or both correct and error trials. Plotting conventions follow what was previously shown in Figure 2B. (C) Quantification of neuronal response dimensionality in exclusively correct trials compared to all trials (correct + error). Data from the two stimulus and the two delay periods are combined.

On the other hand, we reasoned that if increased rotation of neural presentation is relevant for behavior, absence (or reduction) in this rotation will result in more error trials. We thus compared the subspace rotation between correct trials in the Remember 1^st^ context and correct and error trials in the Remember 2^nd^ context. Indeed, we saw that in PFC (Fig. 6B left), the rotation was significantly lower in three out of four trial epochs (1^st^ stimulus,1^st^ delay, 2^nd^ stimulus) when errors are included compared to correct trials alone. The rotation angle for the 1^st^ stimulus epoch decreased from 32.9° ± 3.1° in correct trials, to 25.5° ± 2.2° (*p* < 0.001, permutation test) after including error trials. Similarly, rotation angles decreased from 45.3° ± 3.9° to 40.9° ± 2.7° (*p* = 0.024, permutation test) for the 1^st^ delay, and from 76.4° ± 10.1° to 57.1° ± 14.0° (*p* = 0.042, permutation test) for the 2^nd^ delay epoch. This finding aligns with our RNN modeling results that modest rotation changes in the first three epochs are particularly important for achieving high performance under the Remember 1^st^ context (Fig. 5C left). However, in PPC we did not observe significant rotation changes in error trials. This result also suggests a smaller influence of PPC representations in behavior.

Motivated by these findings, we quantified the dimensionality of neural responses using the participation ratio (Fig. 6C; see Methods for implementation details). We first discovered that during the delay period, neural responses in PFC had significantly lower dimensions than those in PPC (*p* < 0.001, permutation test) consistent with the Overwrite model being more evident in PFC than PPC. Additionally, we found that both PFC and PPC operate in a significantly higher-dimensional subspace in correct than error trials (*p* < 0.001, permutation test). This finding aligns with the observed increase in subspace rotation during both the stimulus and delay periods during RNN training, because larger rotations across time imply the engagement of additional dimensions throughout the trial.

### Nucleus Basalis stimulation improves performance by increasing dimensionality

Although the error trial analysis provided some evidence of neural activity patterns that were predictive of behavior, we wished to test the causality of our predictions in a stronger fashion, by inducing perturbations in neural activity, in an independent sample. We thus wondered if external stimulation could induce changes that quantitatively change how PFC performs our current task. We therefore reanalyzed results from our prior Nucleus basalis (NB) stimulation experiments (Fig. 7A), which can improve monkeys’ performance of the Remember 1^st^ - Remember 2^nd^ task by targeting acetylcholine release ^25^. Overall performance improved from 82.6±0.8% to 90.0±2.2% and from 80.8±1.6% to 83.3±1.5% under stimulation, for the two monkeys respectively. We analyzed a total of 61 neurons recorded from the dorsolateral prefrontal cortex in this study, which incorporated all conditions in both NB stimulation and control (without stimulation) groups.

**Figure 7.**
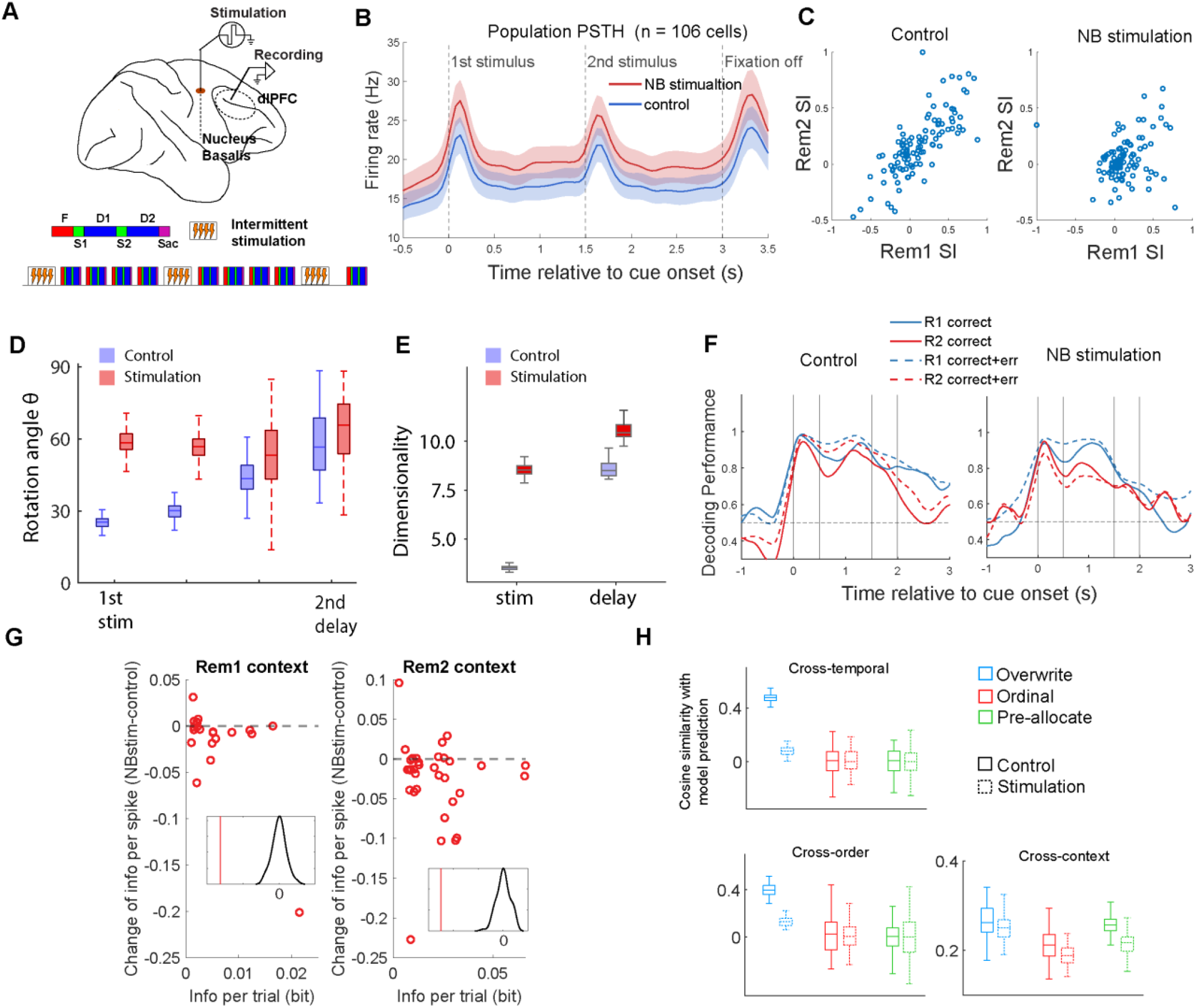
Deep brain stimulation enhances subspace rotation and increases neural response dimensionality. (A) Experimental design for nucleus basalis stimulation. Intermittent electrical stimulation was applied to the nucleus basalis while single-unit recordings were simultaneously conducted in the prefrontal cortex during the context-dependent working memory task. Stimulation was applied during the inter-trial interval, every few trials. (B) Mean firing rate during the control and NB stimulation conditions (n=106 neurons). (C) Single-cell selectivity during the first delay epoch for the two spatial locations was quantified using the Selectivity Index, SI = (R1 - R2) / (R1 + R2), in both the Remember 1st and Remember 2nd contexts. NB stimulation alters single-cell selectivity by rendering neuronal responses context dependent. (D) Mean subspace rotation for four consecutive task epochs: first stimulus presentation, first delay period, second stimulus presentation, and second delay period, under control and NB-stimulation conditions. Deep brain stimulation induces greater subspace rotation. (E) Mean dimensionality during the two stimulus and two delay period under control and NB-stimulation conditions. Deep brain stimulation induces higher dimensionality, quantified by the participation ratio of neural responses. **(F)** Analysis of decoding performance in correct and error trials in the Remember 1^st^ context, where most errors occur. Decoding performance for the location of the first stimulus is shown for both contexts under control and stimulation conditions. In control trials, errors were characterized by reduced context-dependent gating of information after the second stimulus, whereas in stimulation trials, errors were characterized by early differentiation of information. **(G)** Change in the mean information conveyed by each spike in the NB-stimulation relative to the control condition. Red vertical lines in the insets indicate the mean information change, whereas the black curves represent the null distribution obtained by label shuffling. Under NB stimulation, less context-relevant information was conveyed by each spike. (H) Cosine similarity between empirical decoding results and the predictions of the ordinal, pre-allocate, and overwrite models.

To explore potential neural support for this improvement, We first quantified the effects of NB stimulation at the single-cell level. NB stimulation increased the average firing rates of recorded neurons throughout the trial (Fig. 7B). Interestingly, stimulation also altered neuronal selectivity across the two contexts. During the first delay, most cells showed similar preferences in both contexts under control conditions. Under NB stimulation, however, some cells exhibited mixed selectivity, with different preferences depending on the context (Fig.7C). These changes at the single-cell level are directly relevant to the population-level measures analyzed above. For instance, increased mixed selectivity during particular epochs is generally associated with higher dimensionality, whereas elevated firing rates may reflect reduced coding efficiency.

We therefore proceeded to re-examine our population-level measures in this dataset. We first quantified the subspace rotation between the Remember 1^st^ and Remember 2^nd^ context and compared the NB stimulation and control trials (Fig. 7D). The control dataset replicated faithfully the findings of the same analysis in our original dataset: the delay periods had higher rotation angles than the corresponding stimulus periods; and the second delay period which distinguishes the target with the distractor exhibited the highest rotation. Importantly, we discovered that the NB stimulation group had significantly higher rotation angles for 1^st^ stim and 1^st^ delay periods (1^st^ stim *p* < 0.001, 1^st^ delay *p* = 0.03, permutation test). This result is the mirror image of the analysis of error trials (Fig. 6B) and replicates the finding of the RNN analysis (Fig. 5A). We again calculated and plotted the dimensionality under control and NB-stimulation conditions (Fig. 7E). Overall, NB stimulation significantly increased dimensionality for both the stimulus and delay periods (*p* < 0.001, permutation test). Furthermore, when error trials were included in the decoding analysis, we again identified the characteristic signature of the overwrite model, namely reduced context-dependent gating, in the control condition. In contrast, under stimulation, correct trials showed little context modulation of information, whereas error trials exhibited very early context modulation (Fig. 7F). This pattern suggests a possible but unsuccessful attempt to implement an “only remember relevant” strategy. Also, in line with our hypothesis that enhanced performance may come at the cost of greater energy expenditure, we directly quantified the amount of task-relevant information conveyed per spike—specifically, information about the first stimulus location in the Remember 1^st^ context and the second stimulus location in the Remember 2^nd^ context. This analysis showed that the amount of “useful” information per spike decreased under stimulation (Fig. 7G). We finally examined the similarity of stimulus representation under NB stimulation in the Overwrite, Ordinal, and Pre-allocate models. Aligned with rotation and dimensionality results, the similarity of the NB-stimulation condition deviated considerably from the Overwrite model (Fig. 7H), supporting the idea that improved performance conflicts with efficiency that the Overwrite model represents.

## DISCUSSION

Our results demonstrate that the prefrontal and, to a lesser extent, the posterior parietal cortex are capable of efficiently manipulating information that must be maintained during context-dependent working memory tasks. Efficiency arises from encoding task-relevant neural representations into a low-dimensional space that is preserved across time and across different contextual demands. Such low-dimensional structure allows for compact and reusable representations, reducing the computational burden associated with maintaining and transforming information. In contrast, activity in the PPC was more closely aligned with processes responsible for gating or selecting relevant stimulus information, rather than compressing or transforming it. Richly trained recurrent neural networks replicated the representational geometry observed in prefrontal cortical activity, reinforcing the idea that low-dimensional encoding strategies reflect an optimal computational solution under certain constraints. However, efficiency is not without cost. The same compressed representations that allowed for flexible reuse also increased susceptibility to interference and errors, as revealed by analysis of error trials in the empirical data, and RNN simulations. Supporting this interpretation, intermittent electrical stimulation of the Nucleus Basalis, which enhanced behavioral performance, shifted prefrontal activity away from this default low-dimensional regime into a higher-dimensional representational, and thus less efficient, mode. Our findings suggest that PFC prioritizes efficiency but can achieve greater fidelity through transition into a more robust, albeit more resource-intensive, representation. Our results highlight a fundamental tradeoff between efficiency and effectiveness in neural computation.

### A tradeoff between efficiency and effectiveness

Recent studies have demonstrated that low-dimensional representational geometry can provide important insights into the strategies employed by subjects^26^. Using this approach, the current study identified two operational modes of the prefrontal cortex within the same working memory task, which could be reversibly switched by NB stimulation. Recurrent neural network modeling suggested that these modes correspond to distinct learning regimes and network dimensionalities. Although the effects of NB stimulation did not fully conform to either of the two conceptual models we proposed, they nevertheless reproduced several key predictions derived from those models, including increased dimensionality and decreased efficiency. These findings may indicate the existence of a continuous axis governing the trade-off between energetic cost and computational effectiveness. Given that the nucleus basalis of Meynert is the major basal forebrain cholinergic source to the neocortex ^27^, this result suggests that cholinergic modulation may play an important role in regulating this balance. The intuition of the proposed efficiency and effectiveness balance is that a compact, efficient overwrite representation might conserve metabolic resources during information encoding and decoding. However, in a highly recurrent network like the PFC, incomplete erasure of prior memory traces can lead to errors^28^. In contrast, storing information in separate subspaces offers flexibility but may require more resources for encoding and decoding. Even with fully multiplexed coding within the same population, the coding capacity is constrained by population variance compared to independent population coding ^29^. Thus, the findings in current study support the notion that neural efficiency entails costs.

RNNs have increasingly become a powerful tool for inferring the computations underlying neural activity^30, 31, 32, 33^. Representational geometries of neural and RNN activity have been used to infer the behavioral strategies used to solve a cognitive task, which may differ systematically between individuals or species ^26, 34^. Moreover, examining how RNN activity evolves during training can reveal fundamental principles of learning ^35, 36^. The present study extends this framework by proposing that the tradeoff between efficiency and effectiveness may itself be a key factor shaping strategy selection, and that this tradeoff can be quantitatively assessed through analysis of RNN dynamics.

### Prefrontal cortical flexibility

Prefrontal neurons are well known for their ability to represent complex, task-dependent information, often exhibiting mixed selectivity that depends on both stimulus features and contextual demands^2, 37, 38^. Information about distracting stimuli, that subjects recognize as being irrelevant for the task performed, is not always discarded^39^. Instead, such information can be maintained in transformed or orthogonalized representations, allowing the prefrontal cortex to minimize interference ^6, 7, 11^. Additionally, neural resources can be dynamically allocated across sequential stimuli, with individual neurons flexibly participating in representations of multiple sequential stimuli^10^. Against this backdrop, our findings reveal an alternative operational mode in the PFC that emerges when task demands are relatively simple. In our task, which required only a single mental operation across contexts and involved maintaining just one stimulus at a time, the PFC reused the same coding space across conditions. An “Overwrite” model best characterized the neural data in our subjects, suggesting efficient neural resource utilization.

We should note that context cueing in our task was achieved through the color of the fixation point. Individual neurons, particularly in the ventral prefrontal cortex exhibit selectivity for stimulus color ^40, 41^ and in that sense, neuronal responses we recorded may have been influenced by the fixation point color. However, the critical comparison was between the first and second stimulus presentation under the different context, while the fixation point was present throughout the trial.

Diverse representational changes and qualitatively different trained representations across networks initialized with different weights indicate that networks can adopt a continuum of strategies to learn and perform a task. This finding suggests that animal subjects may also rely on different learning strategies depending on their prior experience and underlying circuit architecture. Our animal subjects were trained on single spatial stimulus ocular motor task before trained for the context dependent working memory task, it may be most efficient to adapt the existing neural circuit to use overwrite strategy for a new task. Nucleus basalis stimulation experiments further illuminated the PFC’s adaptability. NB stimulation induced the formation of new, context-specific coding spaces. These results demonstrate the efficacy of NB stimulation and underscore the remarkable flexibility of PFC circuits in supporting working memory.

### Distinct Roles of PFC and PPC

Working memory depends on a distributed network of cortical and subcortical regions, prominently including the prefrontal and posterior parietal cortices^7, 42, 43^. Classical studies have suggested that neurons in the posterior parietal cortex primarily encode the most recent stimulus, particularly in spatial working memory tasks, and that these representations are susceptible to disruption by intervening distractors^44^. In contrast, prefrontal cortex activity has been associated with the maintenance of behaviorally relevant information, even in the presence of distractors^45, 46, 47^. Similar distinctions have been observed in object working memory, where inferior temporal cortex represents recently viewed stimuli, while prefrontal cortex maintains task-relevant information across delays^48^. More recent findings somewhat complicate this picture, showing that prefrontal neurons can, in some cases, respond more strongly to distractors than to the maintained stimulus^49, 50^. Despite these nuances, lesion and inactivation studies suggest that posterior parietal activity generally has a more limited influence on behavior compared to prefrontal cortex^51, 52, 53^.

In recent years, it has become increasingly clear that relying solely on average neuronal firing rates offers a limited view of how brain regions encode information and that single-neuron measures can overlook important dynamics that emerge only when considering activity across populations ^54, 55^. Our current results demonstrate differences in population activity between prefrontal and posterior parietal cortex that may not have been evident in single-neuron activity. First, the posterior parietal cortex showed a greater departure from the predicted efficient “Overwrite” model, indicating that its activity may retain or process information in a more complex or less streamlined manner (Fig. 2E). Secondly, it was possible to decode contextual information about whether the task required remembering the first or second stimulus (Fig. 2F), suggesting that the posterior parietal cortex plays a targeted role in controlling or “gating” incoming information at that stage. Finally, the ability to distinguish between different stimuli during key time periods was weaker in the posterior parietal cortex compared to the prefrontal cortex (Fig. 3E). Taken together, these observations point to distinct functional contributions of the two regions, suggesting that the prefrontal cortex exhibits greater flexibility and adaptability in adjusting to varying task demands and contextual requirements, whereas the posterior parietal cortex may serve more specialized or constrained roles in processing and regulating sensory information.

## METHODS

Data were obtained from two male rhesus monkeys (*Macaca mulatta*), weighing 7–9 kg. Monkeys were either single-housed or pair-housed in communal rooms with sensory interactions with other monkeys. All experimental procedures followed guidelines set by the U.S. Public Health Service Policy on Humane Care and Use of Laboratory Animals and the National Research Council’s Guide for the Care and Use of Laboratory Animals and were reviewed and approved by the Wake Forest University Institutional Animal Care and Use Committee^25, 49^.

### Experimental setup

Monkeys sat with their heads fixed in a primate chair while viewing a monitor positioned 60 cm away from their eyes with dim ambient illumination and were required to fixate on a 0.2° white square appearing in the center of the screen. In order to receive a liquid reward (typically fruit juice), the animals had to maintain fixation on the square while visual stimuli were presented at peripheral locations. Any break of fixation immediately terminated the trial, and no reward was given. Eye position was monitored throughout the trial using a non-invasive, infrared eye position scanning system (model RK-716; ISCAN, Burlington, MA). The system achieved a < 0.3° resolution around the center of vision. Eye position was sampled at 240 Hz, digitized and recorded. The visual stimulus display, monitoring of eye position, and synchronization of stimuli with neurophysiological data were performed with in-house software implemented on the MATLAB environment (Mathworks, Natick, MA), utilizing the Psychophysics Toolbox^56^.

### Behavioral task

The task involved two stimuli appearing in sequence, requiring the monkey to make an eye movement to the remembered location of either the first or the second depending on the color of fixation point (Fig. 1). The animals maintained fixation for 1 second (s), and then two white squares (1.5° in size) were displayed sequentially for 0.5 s, with a 1.5 s intervening delay period between them. The first stimulus (cue) was displayed pseudo-randomly at one of two locations (0° or 180°) or no stimulus. The second stimulus location was either the same as the first, or was displaced by an angle of 45°, 90°, or 180°, so it could be located at 0°, 180°, 225°, 270°, and 315°. There is also a final condition involving no second stimulus presentation. After a second delay period of 1.5s, the monkeys were required to saccade to the location of the first stimulus if the fixation point was white in color (remember-first condition), and to the location of the second stimulus if the fixation point was blue (remember-second condition). In each daily session the two possible locations of the first stimulus were the same for the Remember 1^st^ (Fig. 2A) and Remember 2^nd^ contexts (Fig. 2B), as were the locations of the second stimulus relative to the first. In NB stimulation experiment, both delays were 1 s instead of 1.5 s.

To minimize the uncertainty about the stimulus to be remembered, the remember-first and remember-second conditions were presented in blocks of trials. The animal was required to perform 10 correct trials of the remember-first task, involving presentation of the two stimuli at each possible combination used in the block. The monkeys were rewarded with fruit juice after making a correct saccade. When running these sessions, the monkey was typically tested first with a single-stimulus Oculomotor Delayed Response task^57^ to map the receptive field of neurons recorded, so as to guide selection of the location of stimuli. However, multiple neurons were recorded simultaneously, and it was not possible to optimize the stimulus location for all neurons in a session. We thus identified, post hoc, the location that elicited a better response compared to its diametric for each neuron. We refer to this as the “preferred location” of the neuron, in a relative sense.

### Surgery and neurophysiology

Two 20 mm diameter recording cylinders were implanted over the dorsolateral prefrontal cortex and the posterior parietal cortex of the same hemisphere in each monkey (Fig. 1). Extracellular activity of single units was recorded using arrays of 2–8 microelectrodes in each cylinder, which were either glass-coated, (250 μm diameter, impedance of 1 MΩ at 1 kHz, Alpha-Omega Engineering, Nazareth, Israel) or epoxylite-coated tungsten microelectrodes (125 or 250 μm diameter, impedance of 4 MΩ at 1 KHz, FHC Bowdoin, ME). Electrodes were advanced individually into the cortex with a microdrive system (EPS drive, Alpha-Omega Engineering, Nazareth, Israel). The electrical signal from each electrode was amplified, band-pass filtered between 500 Hz and 8 kHz, and recorded with a modular data acquisition system of 25 μs resolution (APM system, FHC, Bowdoin, ME). The anatomical location of electrode penetration was determined based on MR imaging of the brain. Neuronal data were collected from areas 8a and 46 of the dorsolateral prefrontal cortex (dlPFC) including both banks of the principal sulcus, and areas 7a and lateral intraparietal area (LIP) of the posterior parietal cortex (PPC) in the lateral bank of the intraparietal sulcus.

### Deep Brain Electrode Implantation and Stimulation

The deep brain stimulation procedure was performed on the same two monkey subjects as the regular remember 1st and remember 2nd task. Specifically, once the head-cap and recording cylinders had been implanted, a second surgery was performed to implant the stimulating electrode. Based on MR imaging, stereotaxic coordinates were obtained for targeted implantation. The animals were implanted unilaterally (one in the left, and one in the right hemisphere) at 8mm lateral,16 mm anterior interaural, and 29 mm below the cortical surface in a vertical penetration. The lateral and anterior coordinates, and depth, were chosen to correspond to the center of the anterior portion of the Nucleus Basalis. A small cylindrical titanium chamber (5-mm inner diameter and 7-mm outer) was mounted on the cranium and chamber was encased in bone cement, in continuity with the existing head-cap. A 26 ga. sharp hypodermic guide tube was lowered and the tip advanced 5 mm below the dura mater. The stimulation electrode was inserted into the guide tube, and a stylus was used to push it to the appropriate depth. The guide tube was then raised while the stylus depth maintained. The stimulation pulses were created by an isolated pulse stimulator (Model2100, AM Systems, Sequim WA), which was controlled by custom programed software, written on the MATLAB platform. Impedances of electrodes were checked monthly during experiments. Intermittent stimulation was applied for 15 s at 80 pulses per second, followed by approximately 45 s with no stimulation. Stimulation was applied in the inter-trial interval, after a trial had completed, and a new trial began after stimulation had elapsed. Stimulation was delivered with biphasic, negative first, unipolar 200 μA pulses with 100 ms per phase, and 80 Hz pulses were delivered for 100 ms.

### Neural data processing

A semi-automated cluster analysis was used, relying on the KlustaKwik algorithm ^58^ which applies principal component analysis of the waveforms to sort recorded spike waveforms into separate units. To ensure a stable firing rate in the analyzed recordings, we identified recordings in which a significant effect of trial sequence was evident at the baseline firing rate (ANOVA, p < 0.05), e.g., due to a neuron disappearing or appearing during a run, as we were collecting data from multiple electrodes. Data from these sessions were truncated so that analysis was only performed on a range of trials with stable firing rate. Less than 10% of neurons were corrected in this way.

### Error trial analysis

To understand difference in processing when monkeys made an error, we analyzed error trials. These were trials that were completed, but in which the monkey erroneously made a saccade to a location other than the stimulus that needed to be remembered, defined as deviating by more than 7 degrees of visual angle from the center of the stimulus location. Trials aborted prematurely, e.g. due to a break in fixation before the saccade, were not used for this analysis. Since error trials were fewer than correct, we relied primarily in comparison of correct trials alone with trials including both correct and errors, as we have done previously ^59^.

### Decoding Analysis

We performed a temporal decoding analyses to examine how neural population activity represents and manipulates the task-relevant information over time. Specifically, neural activity was first binned using a sliding time window of 400 ms with a step size of 100 ms. For each time window, we constructed the feature matrix where each element represented an averaged firing rate during that time window of a group of the first and the second locations. The supporting vector machine (SVM) with linear kernel was trained to decode the stimulus direction using this population activity.

All decoding performance was evaluated using out-of-sample generalized accuracy. Neurons with fewer than two trials per condition were excluded, resulting in 187 neurons from the PFC and 224 from the PPC. For each condition, trials were randomly split into two equal sets: one for training and the other for testing. Pseudo-population responses were constructed separately for the training and test sets by randomly resampling 10 trials per condition. A linear SVM decoder was trained on the training set and evaluated on the test set to obtain the decoding accuracy for a single split. This procedure was repeated 100 times with different random splits, and the final decoding performance was reported as the average accuracy across repetitions.

Cross-temporal, cross-order, and cross-context decoding are employed to reveal the working principal of the Remember 1^st^- Remember 2^nd^ task. Cross-temporal decoding was employed to examine how neural representations generalize across time. For this analysis, a classifier was trained at one time point and tested at all other time points, resulting in a two-dimensional decoding matrix that captures the temporal relationship of coded information. Cross-order decoding was used to assess whether the same spatial location, when presented in different temporal orders, shared a common representational subspace. To evaluate this, we trained the decoder on trials in which the stimulus appeared in the first temporal position (first stimulus and first delay) and tested it on trials where the same location appeared in the second temporal position (second stimulus and second delay). Cross-context decoding examined the generalization of neural representations across task contexts (Remember 1^st^ vs. Remember 2^nd^). The decoder was trained on the trials of one context and tested on trials of the other context. Accordingly, the predicted decoding-accuracy matrices for the Ordinal, Pre-allocate, and Overwrite models were constructed based on the assumption that information at a given position is decodable (Fig. 2B). If the information is assumed to be undecodable, the corresponding matrix entry was set to the chance level (0.5 for the first location and 0.2 for the second location). If the information is assumed to be decodable, the matrix entry was set to 0.95. Together, these three decoding matrices provided a comprehensive map of how spatial information is encoded and generalized across time, stimulus order, and task demands.

Results of cross-temporal, cross-order, and cross task decoding result matrix were visualized as a heat map and compared with model predictions using cosine similarity defined as follows:

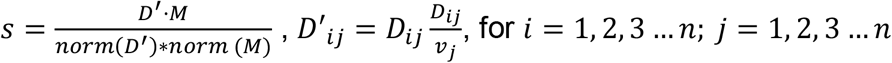

Where *s* represents cosine similarity, *M* is matrix from model prediction, *norm* () indicates 2-norm of a matrix, *D* is matrix for empirical decoding result, *v* is a vector of temporal decoding result representing the amount of decodable information for each time point, and *n* is the number of time points in decoding. In this procedure, raw decoding matrix was first normalized column wise before calculating cosine similarity, to equalize the factor that amount of information varies for each time point.

### Coding neural population activity subspace using principal component analysis

We applied principal component analysis (PCA) to identify a low-dimensional subspace of neural population activity. Previous studies demonstrated that the PCA-based subspace revealed stable population coding for working memory^3^. Specifically, we first calculated the firing rates of each neuron during the first stimulus, first delay, second stimulus, and second delay periods, for two tasks and different stimulus conditions. This resulted in a condition-by-neuron representation matrix. We then normalized the matrix by subtracting the mean of each column to ensure it was zero-centered. PCA was performed on this normalized matrix using singular value decomposition (SVD). We selected the first three eigenvectors, also called the principal components (PCs) of the covariance matrix, sorted in descending order based on the variance explained. These three components defined the subspace of neural representation. In our study, the first three PCs accounted for more than 72% of the variance in the response matrix across all analyzed periods and locations (see Fig. S3), suggesting that neural activity could be effectively represented within the low-dimensional subspace.

### Calculating the subspace rotation between contexts

During each task epoch we defined the corresponding activity matrix for each condition during a given period as an M×N matrix, where M is the number of directions and N is the number of neurons. To find the rotation angle between low-dimensional representations in two task epochs, we concatenated two activity matrices into one single matrix B, with a 2M×N matrix. Then we applied the PCA to project the neural activities of the two conditions into one three-dimensional subspace. After projection, the population representations of the M spatial locations in a specific epoch formed M points represented by an Mx3 matrix.

For the first stimulus and the first delay epochs, there are two directions shown up (M=2). We calculated the rotation between two lines by calculating the cosine angle between two lines. For the second stimulus and the second delay epochs, there are five directions shown up (M=5). We calculated the rotation between two planes. We defined the angles between different conditions based on the best-fit planes for conditions under comparison. Given the vectors spanning the best-fit plane (*P*_1_) for one period are 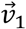 and 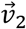 (obtained from the PCs of the reduced eight points), and the best-fit plane (*P*_2_) for another period in the same region are 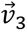 and 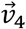, the angle between *P*_1_ and *P*_2_ was calculated as follows:

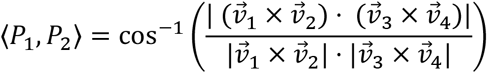

where “⟨*P*_1_, *P*_2_⟩” denotes the angle between *P*_1_ and *P*_2_, cos^−1^ is the inverse cosine function, and 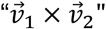 is the cross product that finds the vector that is perpendicular to the plane spanned by 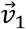 and 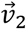, the “⋅” sign stands for the dot product, and “|x|” returns the absolute value (for a scaler) or length (for a vector) of x.

To distinguish between an authentic large rotation of stimulus representation and an incidental rotation arising in a population of neurons with low selectivity and high variability, we calculated each rotation-angle measurement for the control condition. Specifically, we calculated the mean firing rate from n trials for each cell and each stimulus class across different task epochs. For example, we computed the mean firing rate during the first stimulus period from the trials of cell #1 and stored this mean as *λ*. Subsequently, we generated a Poisson distribution with the same mean (*λ*) and randomly produced n sample values from it, simulating n trials from the cue period of this specific cell#1. This control dataset mirrored the firing rate and selectivity statistics of the empirical dataset, thus forming a baseline for rotation within each epoch, and providing a reference for angle and geometry measurements. We assessed variability by randomly drawing 80% of the trials over 100 iterations.

To cross-check the rotation angle between the 2D subspace of different task epochs, we measured the Variance Accounted For (VAF) ratio for each angle measurement1. The VAF ratio for subspace pair (1,2) was defined as follows:

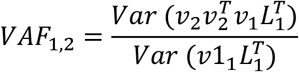

Where *v*_*i*_ (*i* = 1,2) is a 3 × 2 matrix representing two subspace axes, and *L*_1_ represents the stimulus projection in the first subspace. VAF ranges from 0 to 1; large values indicate better alignment of subspaces, while values close to 0 suggest orthogonality.

We implemented a nonparametric bootstrap method to assess the statistical significance of the differences between empirical angles, as well as between empirical angles and control rotation angles. To evaluate whether the inclusion of error trials significantly altered the rotation angle, we performed 1000 bootstrap iterations. In each iteration, we computed the empirical rotation angle difference between 80% randomly sampled correct trials and 80% randomly sampled error trials. This empirical value was then compared to a null distribution of angle differences generated by adding randomly sampled unused trials in the iteration to match the sample size to the dataset including both correct and error trials (named as correct + error in the Results section). The percentage of the empirical value falls within the null distribution was averaged across iterations to obtain a p value. To test whether stimulation affected subspace rotation angles, we performed a similar procedure. In each iteration, we pooled NB stimulation trials with control trials and randomly reassigned them into two groups, preserving the original proportions. A null distribution was constructed from the rotation angle differences between the shuffled groups across 1000 iterations. The p-value was defined as the proportion of times the empirical rotation angle difference fell within this null distribution.

### Multivariable linear regression analysis

To quantify the influence of the first and second stimuli on neural activity during the second delay period, we performed a multivariable linear regression analysis. For each condition, we constructed a one-hot encoded input vector representing the stimulus directions of both the first and second locations. In terms of the first stimuli (0°, 180°, or none) and the second stimuli (0°, 180°, 225°, 270°, 315°, or none), the vector has length of 9 components. For example, if the first stimulus appeared at 0° and the second at 45°, the input vector would be: ***x*** = [1, 0; 0, 1, 0, 0, 0]. The dependent variable ***y***_***i***_ was the average firing rate of the neuron *i* during the second delay period, calculated using the same time windowing as in the temporal decoding analysis. We fit a linear regression model of the form: ***y***_***i***_ = ***β***_***i***_***x*** + ***b***_***i***_, where ***β***_***i***_ is the regression coefficient matrix capturing the influence of each direction (from both the first and second stimuli) on the firing rate, and ***b***_***i***_ is the bias term. For neural activity analysis, due to the limited number of samples, we used lasso regularization to avoid overfitting. The regression coefficient ***β*** ∈ *R*^𝟟×*N*^ allows us to estimate the contribution of each stimulus direction to neural activity at the second delay, which is named as the beta-space representation for both the first and second stimulus locations.

We investigated the neural dynamics by analyzing trajectories in Beta-space and calculated the subspace rotation angles. Specifically, for each context and time period, we obtained a 7×N Beta matrix. To analyze the neural dynamics during the second delay, we concatenated the Beta matrices of all time windows within this period, forming a (7T)×N matrix, where T is the number of time windows. We applied PCA to reduce this matrix to three dimensions, yielding a (7T)×3 matrix, visualized as Fig. 3D.

To assess the rotation angle between the low-dimensional representations of two contexts, we selected Beta vectors corresponding to stimulus directions of interest from each 7×N Beta matrix, forming two M×N Beta matrices for both contexts. These were concatenated into a single 2M×N matrix. PCA was then applied to project this combined matrix into a shared three-dimensional subspace, resulting in a 2M×3 matrix.

### Canonical Correlation Analysis (CCA)

CCA identifies pairs of linear combinations of neural activity, one from each population or condition, that are maximally correlated, thereby quantifying the degree to which two neural subspaces share common representational structure. For each task epoch of interest, we constructed neural population matrices representing the mean firing rate of each neuron across stimulus conditions. Consistent with the rotation angle analysis, conditions for the first stimulus and first delay epochs were defined by the location of the first stimulus, whereas conditions for the second stimulus and second delay epochs were defined by the location of the second stimulus. Prior to CCA, both matrices were mean-centered across samples. CCA was then performed using MATLAB’s built-in “canoncorr” function, which computes the canonical correlation coefficients r_1_, r_2_, …, r_k_ corresponding to k pairs of canonical variates. The primary summary statistic was the mean canonical correlation, defined as: 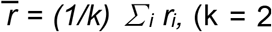, to match the dimensionality of the subspace angle measurement between two planes). To obtain stable estimates of representational similarity and to characterize variability, the full analysis was repeated across 100 independent resamplings of trials. On each iteration, trials were randomly resampled for each condition and neuron, the population matrices were reconstructed, and CCA was performed anew. The mean and standard deviation of 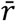 across resamplings were then used to summarize the central tendency and variability of representational similarity for each epoch. Results are reported as mean ± SD across resampling iterations.

### Calculating the location separability

To evaluate how well neural activity patterns encoded spatial information, we quantified location separability, defined as the average silhouette score across stimulus conditions grouped by spatial location. Temporal neural population activity was first projected onto the subspace using the PCA. We then extracted the neural population state at two key time points: early delay (e.g., 0.25s after delay onset) and late delay (1.25s after delay onset). Location separability was defined as the average silhouette score based on predefined labels, the first or second stimulus location. The silhouette score measures how well-separated clusters are by comparing the average distance between points within the same group (same location) to the distance from points in the different group (other different locations). A higher score indicates more distinct clustering of neural representations by location. Silhouette score is defined as follows

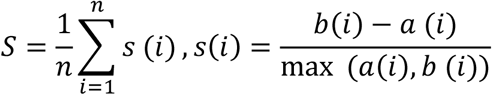

Where S is the overall silhouette score for the dataset, i represent on data point, a (i) represents the average distance from point *i* to all other points within the same cluster (intra-cluster distance), and *b* (*i*) is the smallest average distance from point *i* to all points in any other cluster, excluding its own (nearest-cluster distance). This analysis was repeated over 100 random resampling which randomly samples half of the trials to calculate the firing rate. Paired t-test comparisons between early and late delay periods were performed to assess temporal changes in location encoding.

### Neural Information Analysis

To quantify how NB stimulation altered the efficiency of stimulus-specific neural coding, we computed mutual information between single-neuron firing rates and context-relevant stimulus location (location of the first stimulus under Remember 1^st^ context; location of the second stimulus under Remember 2^nd^ context) in the second-delay epoch. For each neuron, we estimated the mutual information *I* between the discretized firing rate response *R* and the stimulus condition label *S* using the formula:

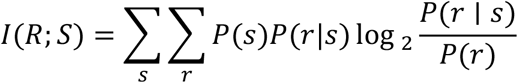

where *P(s)* is the prior probability of stimulus condition *s, P(r)* is the marginal probability of observing firing rate bin *r*, and *P(r*|*s)* is the conditional probability of observing bin *r* given condition *s*. Firing rates were discretized into bins using an adaptive (quantile-based) binning scheme with 5 bins, in which bin edges were defined by the quantiles of the empirical firing rate distribution across all trials for that neuron, ensuring approximately equal occupancy across bins. Information was then normalized by the mean firing rate of the neuron to yield information per spike (bits/spike).

To correct for the positive bias introduced by finite sampling, a shuffle null distribution was constructed for each neuron by randomly permuting condition labels across trials (100 permutations), recomputing mutual information for each permutation, and taking the mean of the resulting distribution as an estimate of the bias (*I*_shuffle). Neurons were included in the change of information analysis if their observed mutual information (bits) exceeded the mean of the corresponding shuffle null distribution (*I*_obs > *I*_shuffle_mean). To assess the statistical significance of the observed information change, we constructed a permutation null distribution by randomly interleaving control and stimulation trials for each neuron and reassigning them to two pseudo-groups of equal size matching the original trial counts. This shuffling procedure was repeated 100 times. For each permutation iteration, information per spike was recomputed for both pseudo-groups across all neurons, and the mean information change was computed using the same selective neuron indices identified from the empirical data.

### RNN modeling and training

We employed recurrent neural networks (RNNs) to investigate the computational mechanisms underlying context-dependent working memory. The network architecture consists of input, recurrent, and output layers, as illustrated in Fig. 4A. Gated recurrent units (GRUs) were used as the basic component of the recurrent layer ^21^. The GRU introduces gating mechanisms that control information flow, allowing the model to retain or discard information dynamically, which helps maintain long-range dependencies in sequences. The dynamical updating functions of the GRU are defined as follows:

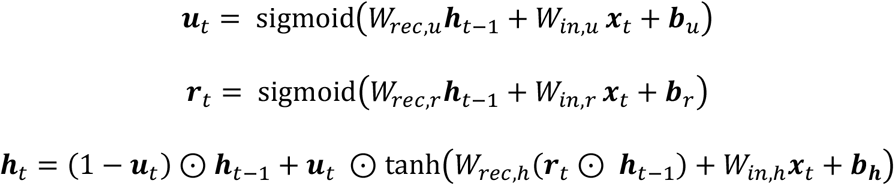

where ***h***_*t*_ is the hidden state vectors. The ***u***_*t*_ term is the update gate that determines how much new information is updated. The ***r***_*t*_ term is the reset gate, determining the extent to which the activity of nodes is used to update the activity of other nodes. The recurrent layer contained N = 128 GRU units. All weights *W*_*rec,u*_, *W*_*rec,r*_, *W*_*rec,h*_ and biases ***b***_*u*_, ***b***_*r*_, ***b***_*h*_ are learnable parameters of the model. The symbol ⊙ denotes the element-wise multiplication. The vector ***x***_*t*_ is the extrinsic input, which encodes the stimuli, fixation, and contexts of Remember 1^st^ or 2^nd 33^, specifically, the inputs including fixation, stimulus, task vectors, and the input noise.

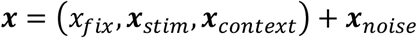

The fixation input is a binary signal indicating whether the subject was fixating (1) or not (0). The stimulus input was encoded using a ring-like structure of eight directionally tuned units that uniformly covered angles from 0 to 2*π*. For a stimulus ***x***_*stim*_ with direction *γ*, the value of the *i* unit with preferred direction *γ*_i_ was defined by:

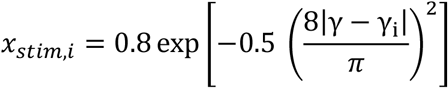

The context input is a one-hot vector with two units corresponding to the Remember 1^st^ and 2^nd^ contexts, respectively. Additive Gaussian noise ***x***_*noise*_ ∼**𝒩**(0, 0.01) was included in the input vector to simulate trial variability. Together, the fixation (1 unit), stimulus (8 units), and context (2 units) components yielded a total of 11 input units. The stimulus direction and temporal structure followed the same design as in the main task (Fig. 1A). Each time step in the model corresponded to 50 ms in real time.

The hidden states ***h***_*t*_ were linearly transformed to the predicted output vector as: 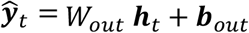. The output 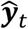 was trained to predict the label ***y***_*t*_, which includes the fixation output *y*_*fix*_, and the target saccadic direction ***y***_*stim*_. The fixation output is greater than 0.85 before the response period and 0.05 during the response period. The target saccadic direction is encoded by the same 8-unit ring, identical to the input encoding. Each unit corresponds to a preferred *γ*_i_ ∈ [0,2*π*), and the target activity is defined as:

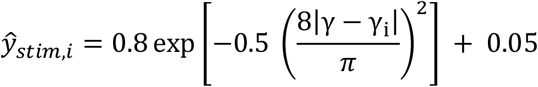

The response direction is read out from the population average response of the 8-unit direction^60^:

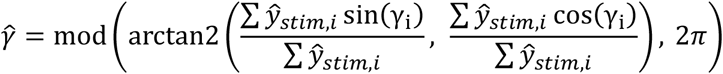

To enhance the long-term dependency and improve the fluent gradient descendent, we additionally applied intermediate supervision at an internal time point^61^. Specifically, after the target direction appears, we introduced an auxiliary supervision term encouraging the hidden state to begin aligning with the saccade direction. We used an auxiliary linear readout layer to predict the target location as: 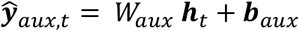. The label ***y***_*aux,t*_ is the same as the target saccadic direction. This supervision mimics brain’s maintenance of the target direction in their persistent activity of the PFC before the response period.

The network was trained to minimize a loss function *L*_*MSE*_, defined as the time-averaged mean squared error (MSE) between the predicted output 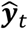 and the target output ***y***_*t*_ across output units i and time points t. The loss is weighted by a non-negative mask matrix *m*_*i,t*_, defined as 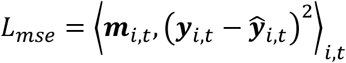. During the stimulus and delay period, the ring output units has the mask weight equal to 1. In the response epoch, the first 100 ms was treated as a grace period with *m*_*i,t*_ = 0, followed by an elevated weight for the response period *m*_*i,t*_ = 5. For the fixation output unit, we used the *m*_*i,t*_ = 2, to prioritize stable fixation control.

All weights were initialized using a normal distribution. The initial input, output, and auxiliary weights (*W*_*in*_, *W*_*o*ut_, *W*_*a*ux_) were drawn from 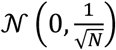, where N = 128 is the dimension of hidden state. The recurrent weights were also initialized by normal distribution 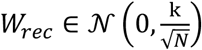, with the weight initialization scaling factor *k* ranging from 0.2 to 3.0 in increments of 0.4. The value of *k* determined whether the models were rich or lazy trained. For each *k*, 20 models were trained by setting the random seed from 0 to 19. All biases are initialized to be zero.

Training was performed using batched gradient descent with a batch size of 128. The Adam optimizer was used with a learning rate of 0.001 and exponential decay rates of 0.9 and 0.999 for the first and second moment estimates, respectively. Model performance was evaluated based on the accuracy of saccadic responses during the response period. A response was considered correct if the decoded direction deviated from the target direction by less than 36 degrees. Each model was trained until it reached 100% accuracy on the task.

### Representational dissimilarity matrix analysis

We utilized representational dissimilarity matrix (RDM) analysis to compare neural response between brain areas and RNNs^24^. The RDM is a condition-by-condition matrix, where each condition is defined by a specific context (R1 or R2), the first stimulus, the second stimulus, and the time epoch. Each element of the RDM corresponds to the Euclidean distance between the average neural firing rates across all recorded neurons for a given condition. This matrix thus captures the similarity of neural population responses across task conditions, providing a representation of the task space. We computed RDMs separately for each brain region and time window, allowing for comparisons of neural representation across phases and areas.

### Training and evaluation of Ordinal and Pre-allocate RNNs

To implement alternative solutions to the Remember 1^st^ –Remember 2^nd^ task, we trained RNNs with the same recurrent architecture and input/output structure used for the Overwrite model, but with modified additional readout heads. Specifically, all models received the same task inputs and were trained on the same context-dependent working memory task. In those trained RNNs in the last section which exhibited Overwrite type representation, RNNs are trained using a single location output that decoded the relevant target representation. By contrast, we added the Ordinal and Pre-allocate models instead with two location output heads, namely Head A and Head B. For the Ordinal model, Head A was trained to reconstruct the first stimulus and Head B to reconstruct the second stimulus. For the Pre-allocate model, Head A was trained to reconstruct the target and Head B the distractor. Besides the training loss for the Overwrite RNNs, we added another orthogonality penalty on the two readout heads to encourage two head encode two orthogonal subspaces^62^

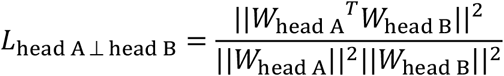

Then we have

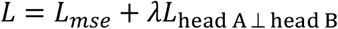

with *λ* = 0.001. We trained the Ordinal and Pre-allocate models until the accuracy reached 100% accuracy on the task and also the *L*_head A ⊥ head B_ < 0.001. All Ordinal and Pre-allocate RNNs are trained under rich training mode (weight initialization scaling factor *k* = 1.0). We used the Overwrite RNNs trained with *k* = 1.0 to compare with them.

After training, we evaluated the models using the same procedures applied to the Overwrite networks. To quantify efficiency, we measured activity energy as the L2 norm of hidden activity. To quantify effectiveness, we re-evaluated trained networks after adding Gaussian noise to the stimulus input, using different noise level (the standard deviations from 0.01 to 0.09) and measured task accuracy separately for the Remember 1^st^ and Remember 2^nd^ conditions across increasing noise levels.

### Neural Representational Dimensionality

We quantified the representational dimensionality of neural activity using the participation ratio (PR) ^63^. We calculated the average neural response for each condition and each time epoch. The representational dimensionality of the average neural response was estimated using the participation ratio, defined as:

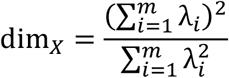

where dim_*X*_ denotes the dimensionality of region *X*, and *λ*_*i*_ represents the *i*-th eigenvalue of the neural representation of region *X*, with a total of *m* eigenvalues. This measure reflects the effective number of dimensions that contribute to the variance of the representational space. A flatter eigenvalue spectrum (i.e., more uniform distribution of variance across components) results in a higher participation ratio, indicating higher dimensionality. Conversely, when a small number of eigenvectors explain most of the variance, the dimensionality is lower.

## ACKNOWLEDGEMENTS

Supported by NIH award number R01 EY036089. We wish to thank Chrissy Suell and Kayla Yetman for technical help and Zhengyang Wang, Rana Mozumder and Alireza Karimi, for helpful comments on an earlier version of this manuscript.

## AUTHOR CONTRIBUTIONS

CC designed the experiment. WD and PC performed data analyses. PC performed modeling. CC, WH, and PC wrote the paper.

## DATA AVAILABILITY

Data for the current study will be made available through Zenodo upon acceptance of the paper

## CODE AVAILABILITY

The code used to process the results and generate the figures will be made available at CodeOcean upon acceptance of the paper

## Supplementary Information

### SUPPLEMENTARY TEXT

#### Mathematical proof formalizing the tradeoff between efficiency and cost

A recent theoretical framework by Dorrell et al.^16^ utilized rigorous mathematical deduction to demonstrate that, under constraints of metabolic or computational efficiency, the subspace alignment angle is determined by the correlation between stimulus orders. Specifically, stimuli with higher correlations are predicted to have smaller angles between their respective coding subspaces, while independent stimuli should occupy orthogonal subspaces. Our empirical findings align with this prediction, as the stimuli presented across the two task epochs were not perfectly decorrelated. Conceptually, an “overwrite” model is more efficient because it reduces the total information load; however, it may be less advantageous due to potential interference caused by the incomplete erasure of task-irrelevant information. We adapted the framework of Dorrell et al. to formally test whether: 1) the “overwrite” model maintains a lower minimal energy requirement compared to “store-both” models; 2) the “overwrite” model exhibits decreased effectiveness, characterized by increased decoding error under noise due to leakage or cross-epoch interference.

Consider a single delay representation *r* ∈ ℝ^*N*^ that must allow linear decoding (with decoding matrix *U* ∈ ℝ^*N*×*m*^) of some task variable *z* ∈ ℝ^*m*^ with zero error:

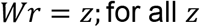

Assume we encode *z* (with encoding matrix *U* ∈ ℝ^*N*×*m*^) linearly:

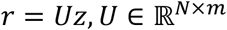

Then the exact-decoding constraint becomes

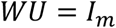

If *r* = *Uz* and *z*’s covariance is Σ_*z*_ = 𝔼[*zz*^⊤^], then

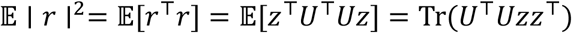

If we whiten the task variables i.e. Σ_*z*_ = *I*, then

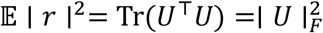

Following Dorrell et al., we define efficiency as an optimization problem to minimize the sum of neural activity and dendritic weight:

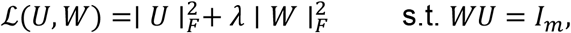

Where *m* is the dimension of the variable you want to reconstruct. If *N* is the number of neurons, and *λ* > 0 is a tradeoff hyperparameter determining the relative contribution of neural activity and weight to energy cost, it could be proven (see below) that:

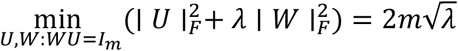

Specifically, to solve this constrained minimization

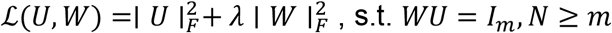

We fix any feasible U, and consider the inner problem:

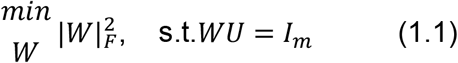

The unique minimizer is the Moore–Penrose left pseudoinverse

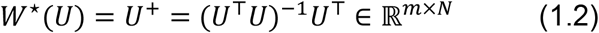

And its Frobenius norm is

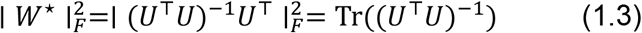

So, problem (P) reduces to minimizing over *U* only:

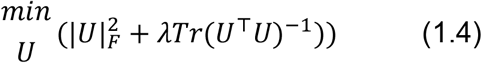

Taking the SVD:

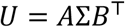

where *A* ∈ ℝ^*N*×*m*^ with *A*^⊤^*A* = *I*_*m*_, *B* ∈ ℝ^*m*×*m*^ orthogonal: *B*^⊤^*B* = *I*_*m*_, Σ = diag(*σ*_1_, …, *σ*_*m*_) with *σ*_*i*_ > 0, then

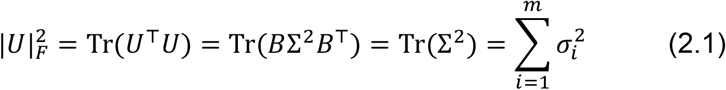

From *U*^⊤^*U* = *B*Σ^2^*B*^⊤^,

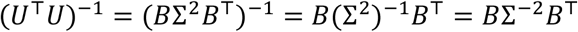

So

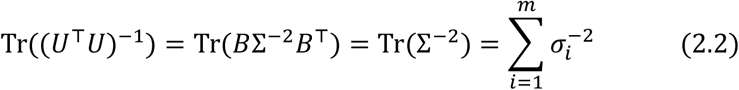

Substituting (2.1) and (2.2) in (1.4) we get

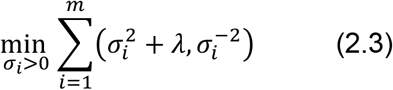

For a single *σ* > 0, define

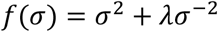

Differentiate:

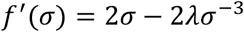

Set *f*^′^(*σ*) = 0:

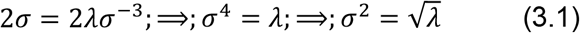

At this *σ*,

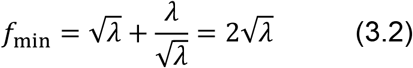

Since the sum in (2.3) is separable across *m*, the minimum is

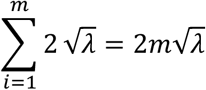

In other words, the minimum cost scales linearly with the dimensionality of the task variables maintained in memory. Furthermore, within this linear framework, it can be seen that the cost also scales with the effective dimensionality (r) of the neural representation.

For the second proposition, we focus on the critical case of the Remember-2^nd^ context, where overwrite must erase the first memory.

Let the ideal overwrite update be “replace *z*_1_ with *z*_2_”. We could write the end-of-trial representation as

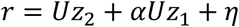

where:*α* is residual leakage of the first stimulus after overwrite (*α* = 0 is perfect overwrite; *α* ≠ 0 is incomplete erasure / imperfect gating), and 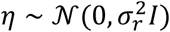 is activity noise. With a decoder *W* satisfying *WU* = *I*, the decoded output is

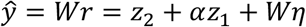

So the decoding error is

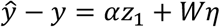

Thus the MSE (effectiveness inverse) is

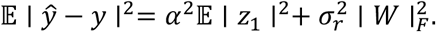

Then we can compare it to store-both (Ordinal or Pre-allocate) models. In Pre-allocate (same conclusion for Ordinal), with ideal subspace separation,

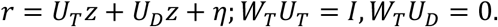

Then

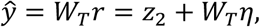

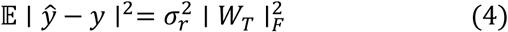

So there is no *α*^2^𝔼 ∣ *z*_1_ ∣^2^ interference term in this case. That means with any nonzero overwrite leakage, the overwrite will be less precise.

The proof above illustrates a cross-modal comparison indicating that while the Overwrite mode offers superior efficiency, its performance may degrade more in the presence of noise. Furthermore, within the same framework, we could also prove a trade-off between efficiency and effectiveness specifically within the Overwrite model itself.

Let the maintained variable be

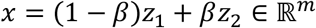

Again, let the neural representation in the second delay under Remember 2nd context be

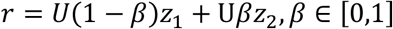

where *U* ∈ ℝ^*n*×*m*^ is the encoding matrix, with *m* ≥ *d*. And β is the leakage parameter

Let *W* ∈ ℝ^*m*×*n*^ be the linear decoder, and impose the decoding constraint

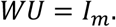

We minimize the same efficiency objective

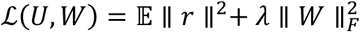

subject to

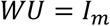

Since *r* = *U*x,

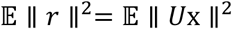

So the optimization problem is

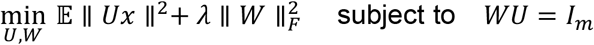

We first compute

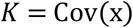

By definition,

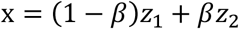

Hence

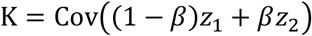

Using the bilinearity of covariance,

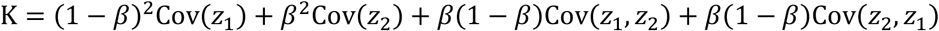

By assumption,

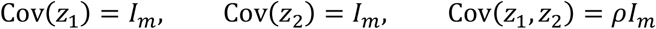

Since *ρI*_*d*_ is symmetric,

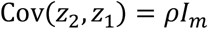

Therefore

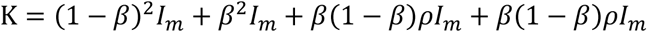

So

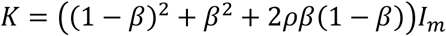

Define

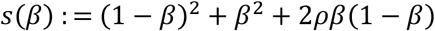

Then

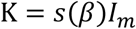

Now compute

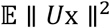

Since ∥ *U*x ∥^2^= x^⊤^*U*^⊤^*U*x,

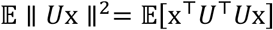

Using the identity 𝔼[x^⊤^*A*x] = tr*A*Cov(*x*x) for zero-mean *x*,

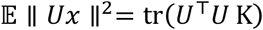

Because *K* = *s*(*β*)*I*_*m*_,

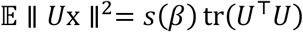

Therefore, the objective becomes

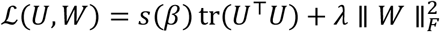

So the optimization problem is

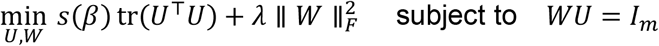

Let the singular values of *U* be

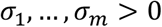

They are positive because *WU* = *I*_*d*_ implies *U* has full column rank *d*.

Since

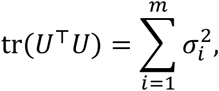

the first term is

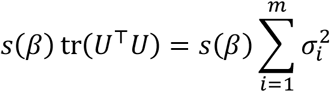

Now minimize over *W* subject to *WU* = *I*_*m*_.

For a fixed full-column-rank *U*, the minimum-Frobenius-norm solution of *WU* = *I*_*m*_ is the Moore-Penrose left inverse

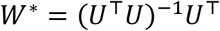

Its singular values are

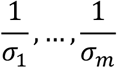

Therefore

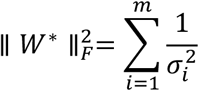

So after minimizing over *W*, the problem reduces to minimizing

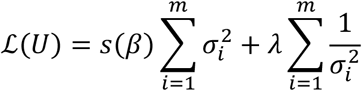

Equivalently,

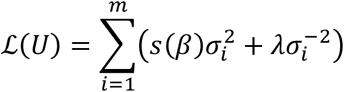

Fix one singular value *σ* > 0. Define

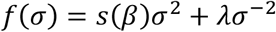

Differentiate:

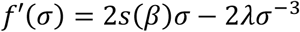

Set the derivative to zero:

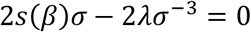

This yields

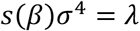

Hence the minimizer satisfies

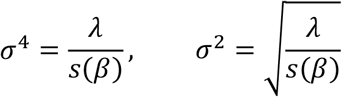

Now evaluate *f* at this minimizer:

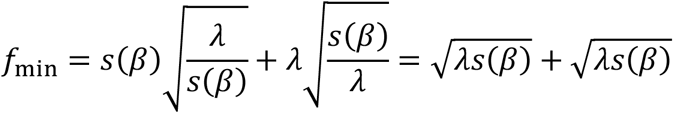

So

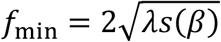

Since there are *m* identical independent terms, the total minimum is

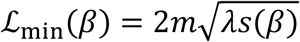

Thus

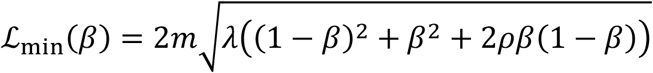

Using

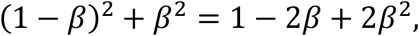

we can also write

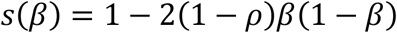

So equivalently,

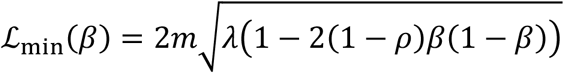

This is the exact minimum efficiency cost under the stated efficiency definition.

Now differentiate *s*(*β*):

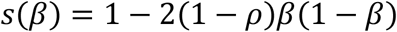

So

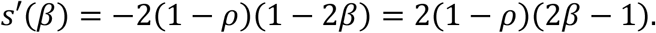

Since 0 ≤ *ρ* < 1, we have 1 − *ρ* > 0. Therefore:

- if β< 1/2, then *s*′(*β*) < 0
- if *β* = 1/2, then *s*′(*β*) = 0
- if *β* > 1/2, then *s*′(*β*) > 0

Because

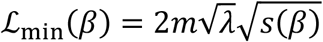

and square root is increasing,

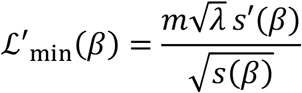

Hence

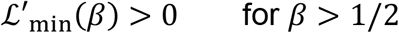

Therefore,

For *β* ∈ [1/2,1](i.e. over chance performance) the minimum efficiency cost increases with overwrite strength.

## SUPPLEMENTARY FIGURES

**Supplementary Figure S1.**
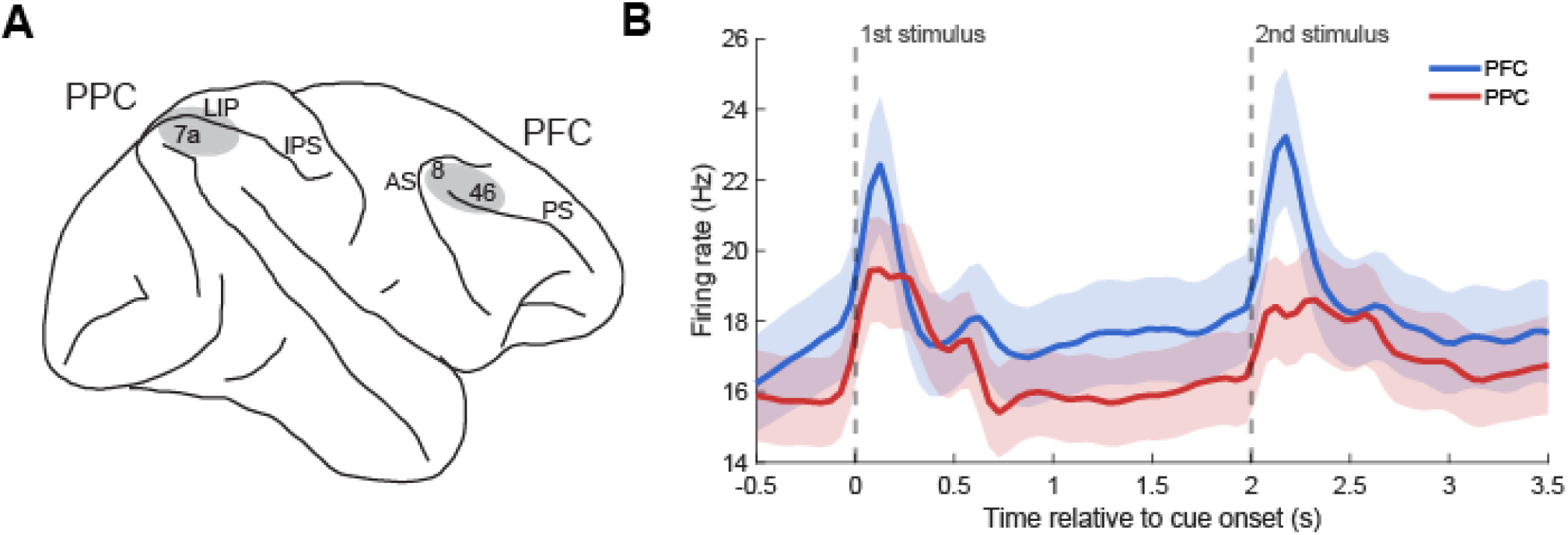
Recording sites and neural responses. (A) Diagram of the monkey brain. Areas that were used for recording in the dorsolateral prefrontal (PFC) and posterior parietal (PPC) cortex are highlighted in gray. Abbreviations: IPS: intraparietal sulcus; LIP: lateral intraparietal area; AS: arcuate sulcus; PS: principal sulcus. (B) Mean firing rate of neurons recorded from the prefrontal and posterior parietal cortex.

**Supplementary Figure S2.**
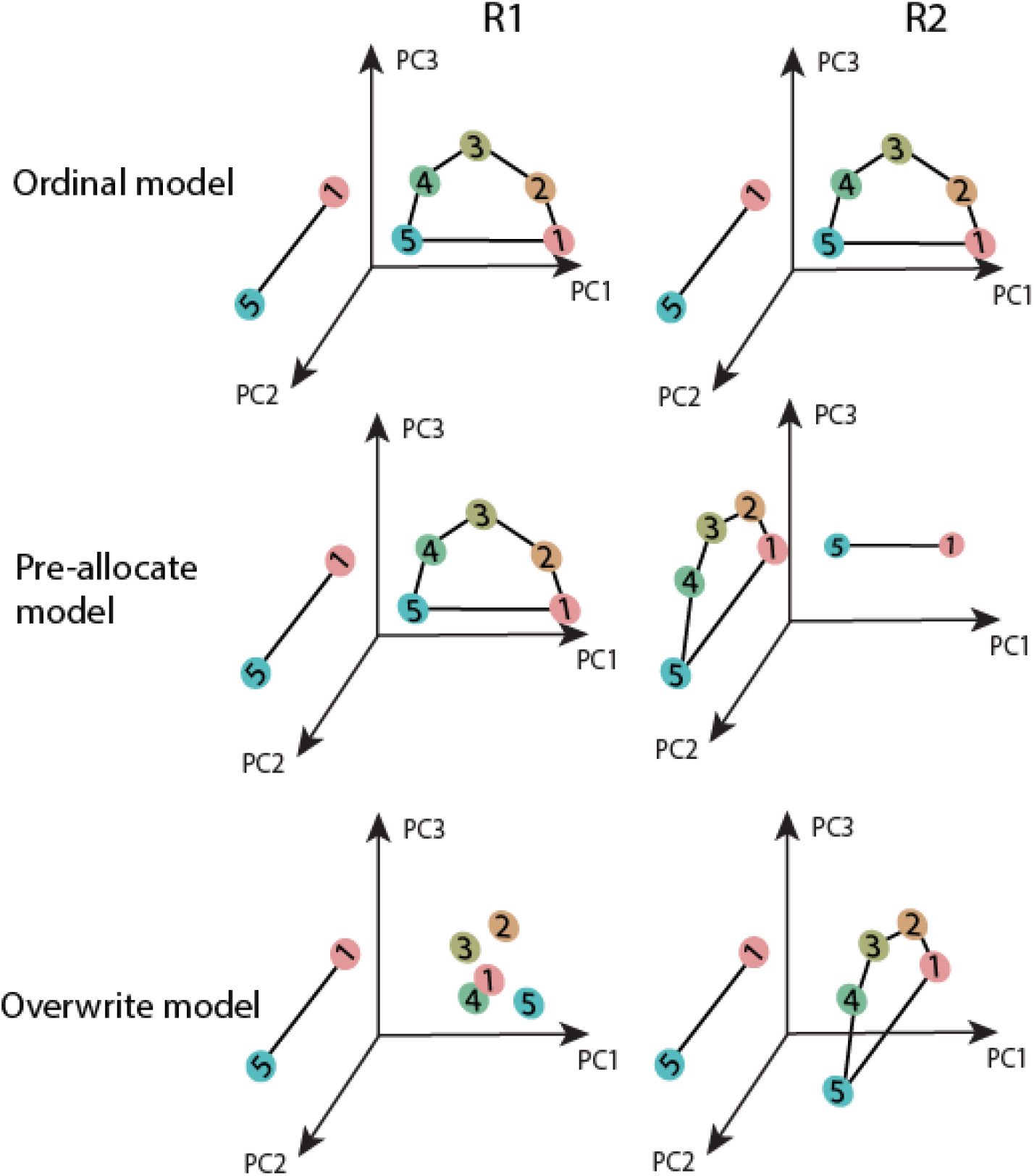
Neural population rotation between contexts. Diagram showing the predicted rotation angles of the ordinal, pre-allocate, and overwrite models. Lines represent the mean neural responses for the two locations presented during the 1^st^ stim and 1^st^ delay epochs. Polygons represent neural responses for each of the 5 locations that were used during the 2^nd^ stim and 2^nd^ delay epochs, averaged across two of the 1^st^ stim epoch locations. Rotation angles were calculated between lines defined by the two stimuli appearing during the 1^st^ stimulus interval and between the planes defined by the five stimuli appearing during the 2^nd^ stimulus interval.

**Supplementary Figure S3.**
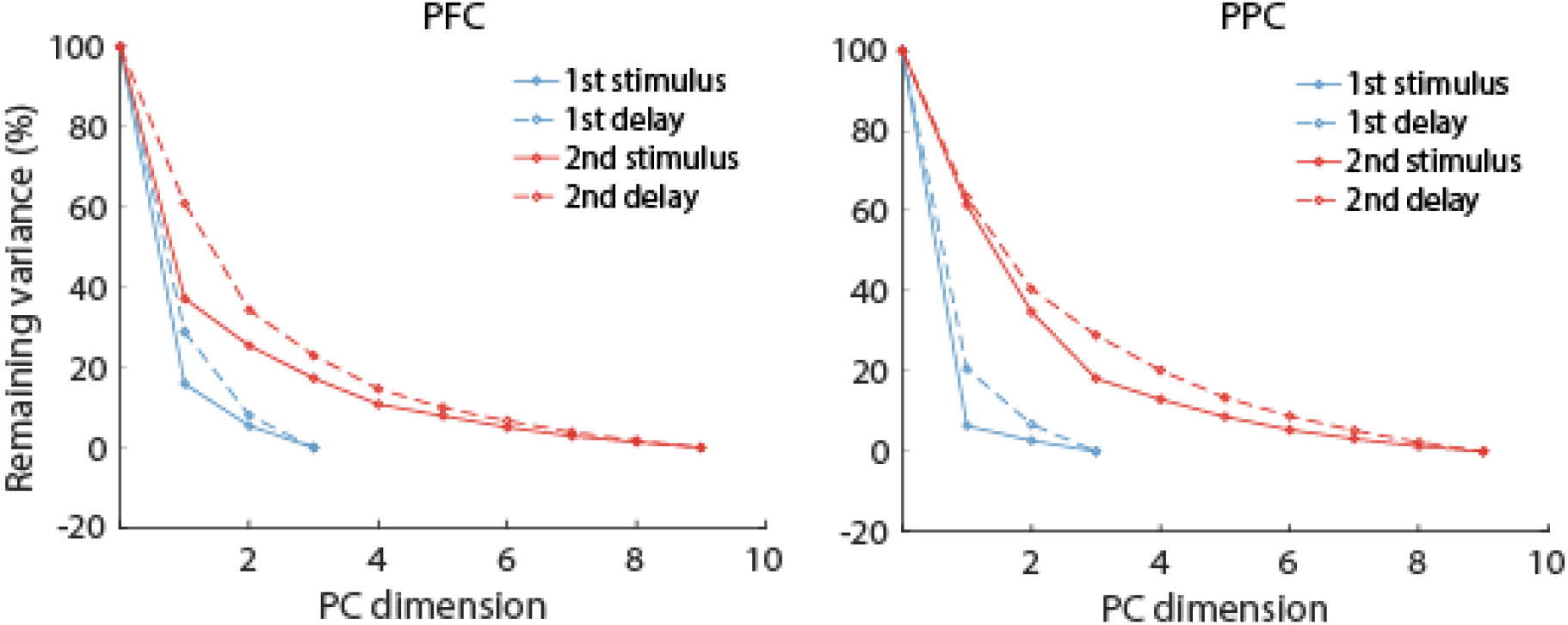
Dimensionality of neural representations. Unexplained variance as a function of increasing principal components for each of the different task epochs for the prefrontal cortex (left) and the posterior parietal cortex (right).

**Supplementary Figure S4.**
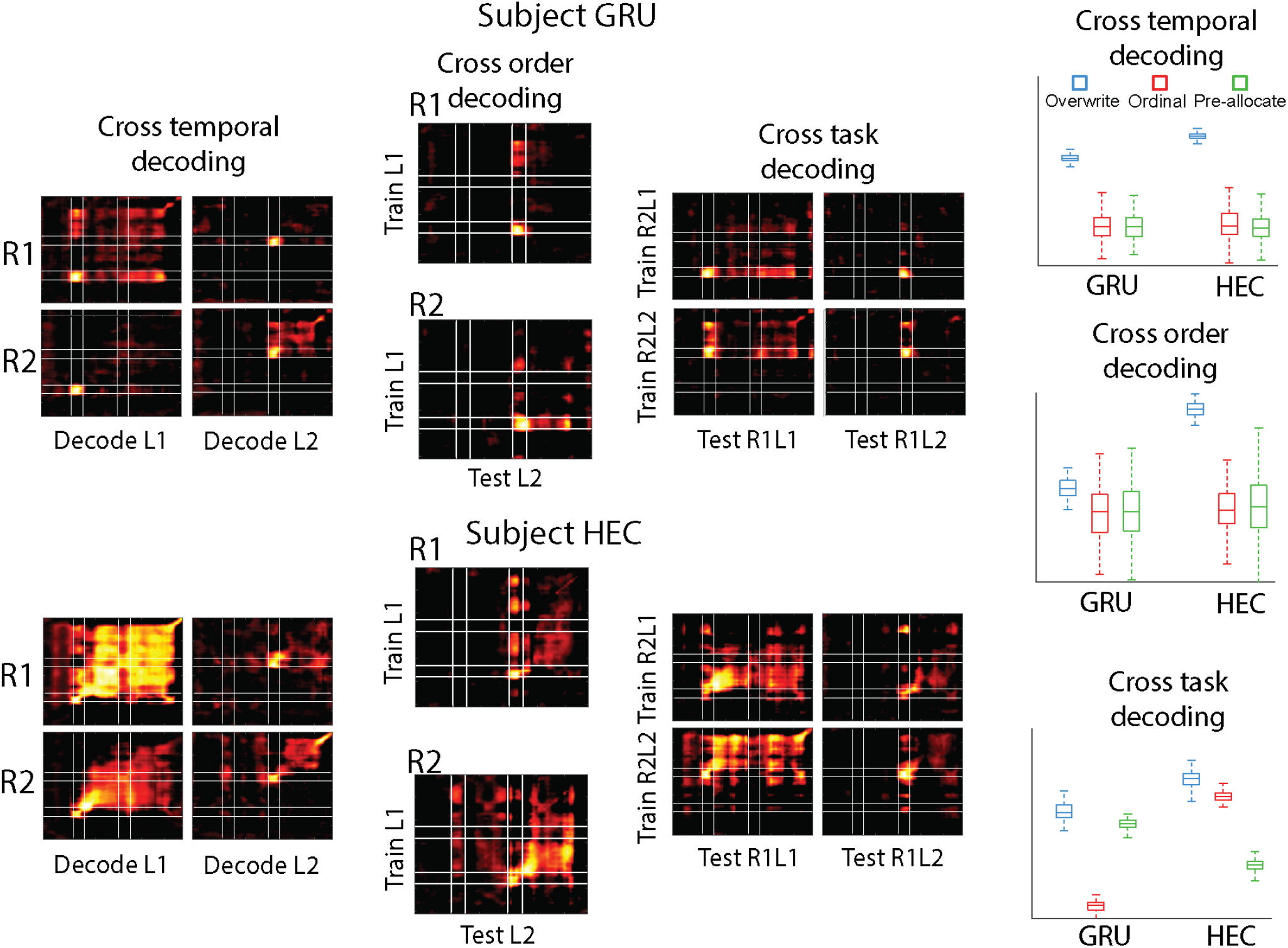
Individual subject decoding. Cross-temporal, cross-order, and cross-task decoding performance for the individual animal subjects (GRU and HEC), shown in the same format as Fig. 2C-E.

**Supplementary Figure S5.**
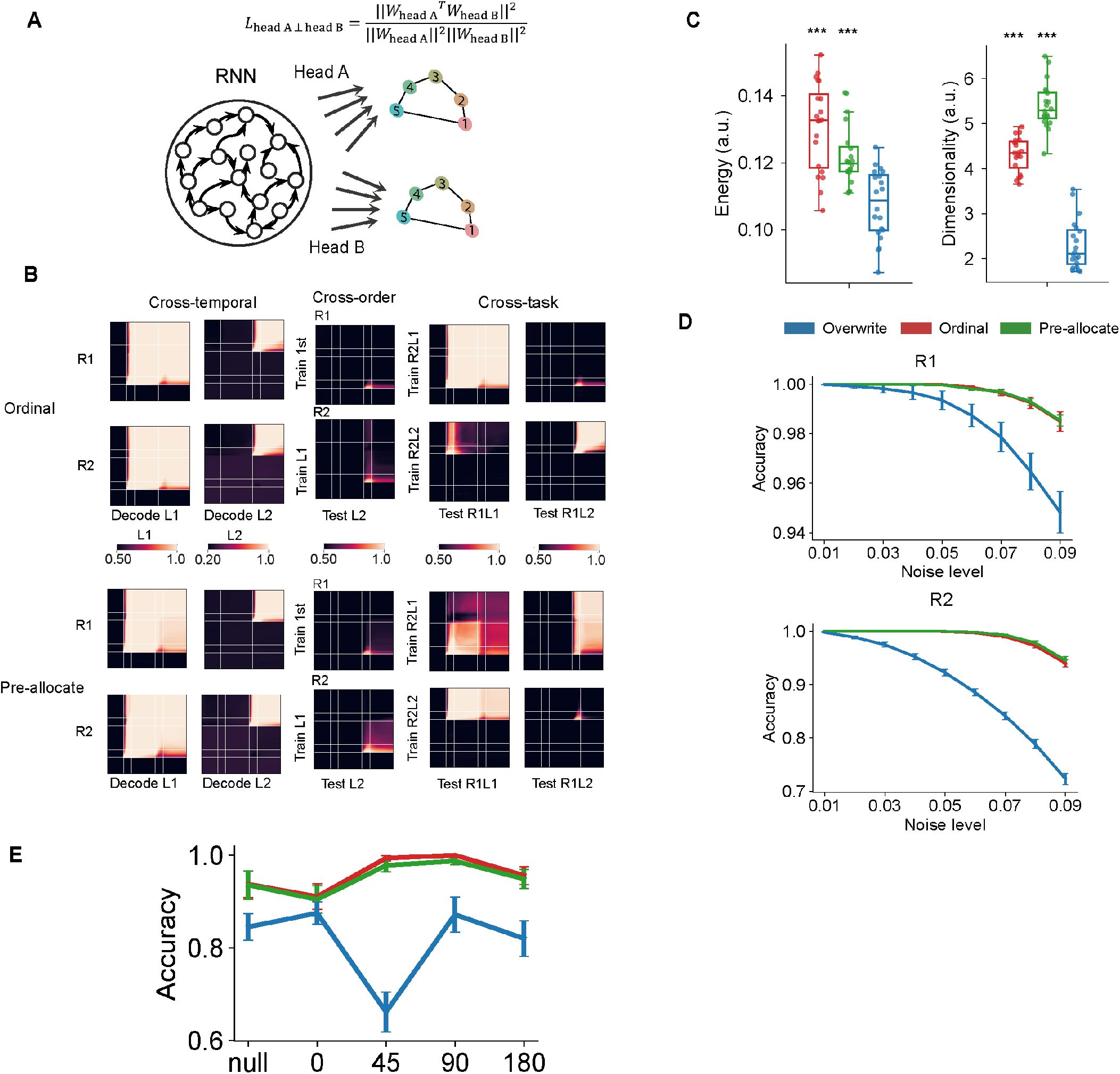
Ordinal and Pre-allocate RNNs are less efficient but more effective than Overwrite RNNs. (A) Schematic illustration of the training structure for the Ordinal and Pre-allocate RNNs. Two Heads (A and B) are trained with an orthogonality loss to maintain two subspaces for two locations. In the Ordinal model, Head A and Head B support the first and second stimuli, respectively. In the Pre-allocate model, Head A and Head B decode target and distractor information, respectively. A sample Remember 2nd trial is shown. (B) Cross-temporal, cross-order, and cross-context decoding matrices for the Ordinal (top) and Pre-allocate (bottom) RNNs. (C) Mean activity energy and effective dimensionality of representations in the Ordinal, Pre-allocate, and Overwrite models. The Ordinal and Pre-allocate models showed significantly higher energy consumption (left) and higher effective dimensionality (right) than the Overwrite model, as indicated by *** (*p* < 0.001, two-tailed t-test). Dots represent individual trained networks. (D) Task accuracy as a function of added stimulus noise for the R1 and R2 tasks across different noise levels (standard deviation of Gaussian noise). Ordinal and Pre-allocate RNNs maintained better performance under perturbation, indicating greater effectiveness. (E) Performance of different RNN models as a function of target-distractor distance.

**Supplementary Figure S6.**
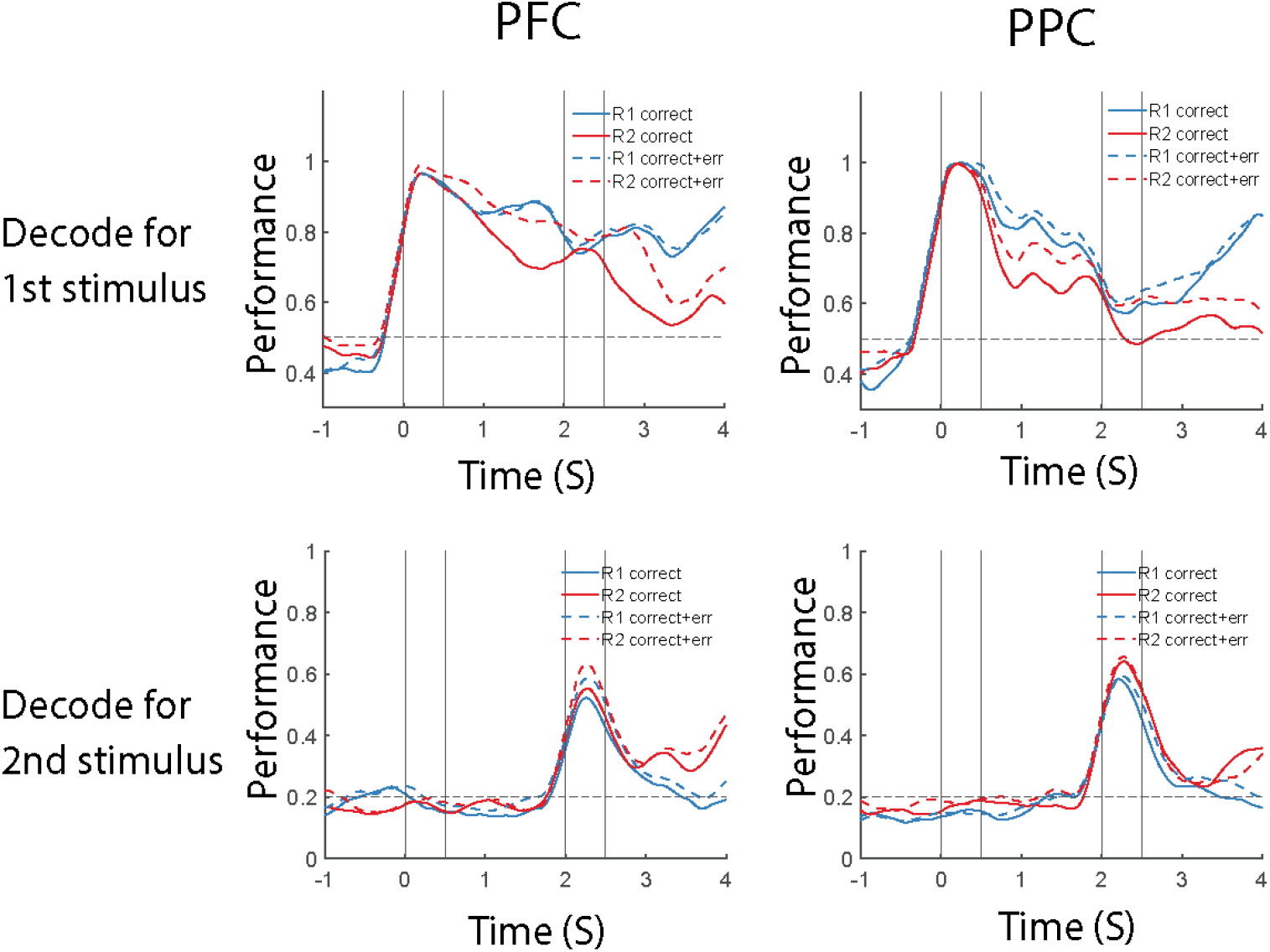
Decoding performance in correct and error trials. Comparison of decoding performance for correct trials versus combined correct and error trials across time is shown for the PFC and PPC.

**Supplementary Figure S7.**
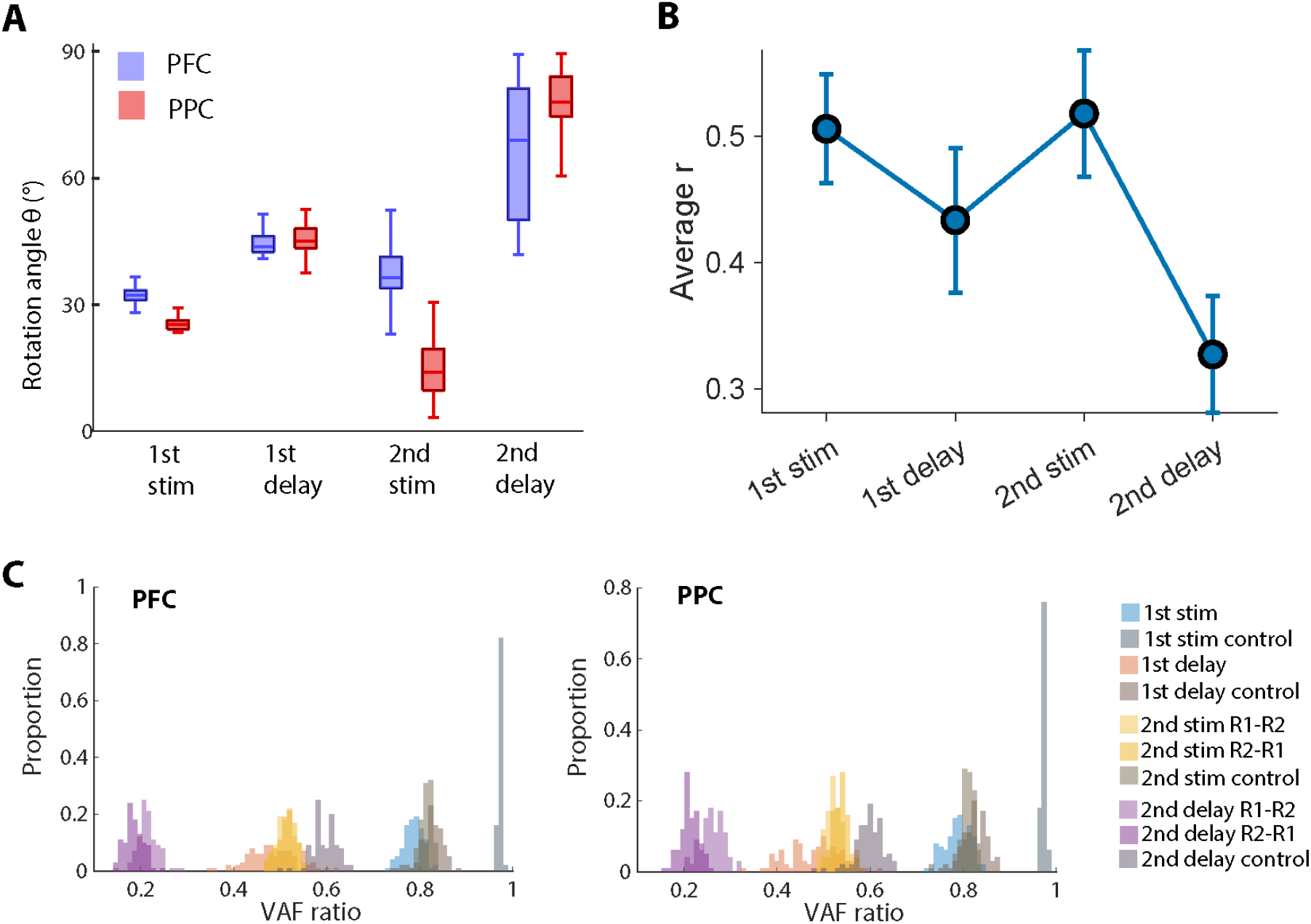
Alternative determination of subspace rotation. (A) Subspace rotation angles between Remember 1st and Remember 2nd contexts across four task epochs, derived by fitting single-cell responses to task variables via lasso regression, followed by principal component analysis on the regression coefficients. (B) Average correlation coefficient (*r*; dimensionality = 2) between contexts, measured using canonical correlation analysis (CCA). The results are quantitatively consistent with the rotation measures, such that smaller angles correspond to higher correlations. (C) Variance accounted for (VAF) ratio between subspaces. VAF measures the degree to which the neural activity in one context can be reconstructed using the principal components derived from a second context. Smaller VAF values indicate higher rotations. Due to the asymmetry of VAF computation, we were able to display the two projection directions (“1-2” and “2-1”) separately by using the representations from the Remember 1st and Remember 2^nd^ contexts, respectively. For the 1st stimulus and delay, only two stimulus directions were used, thus making the VAF computation effectively symmetrical (“1-2” = “2-1”) and resulting in a single distribution. To evaluate whether the observed VAF values reflect meaningful structure, we generated a control distribution via controlled representations constructed for the subspace rotation analysis. These controlled distributions, shown in gray, preserve firing rate statistics but eliminate task-dependent structure. In both the prefrontal cortex and the posterior parietal cortex, the empirical VAF distributions were significantly left-skewed compared to the control (*P* < 0.0001, using non-parametric permutation test), indicating different neural coding for the two contexts.

**Supplementary Figure S8.**
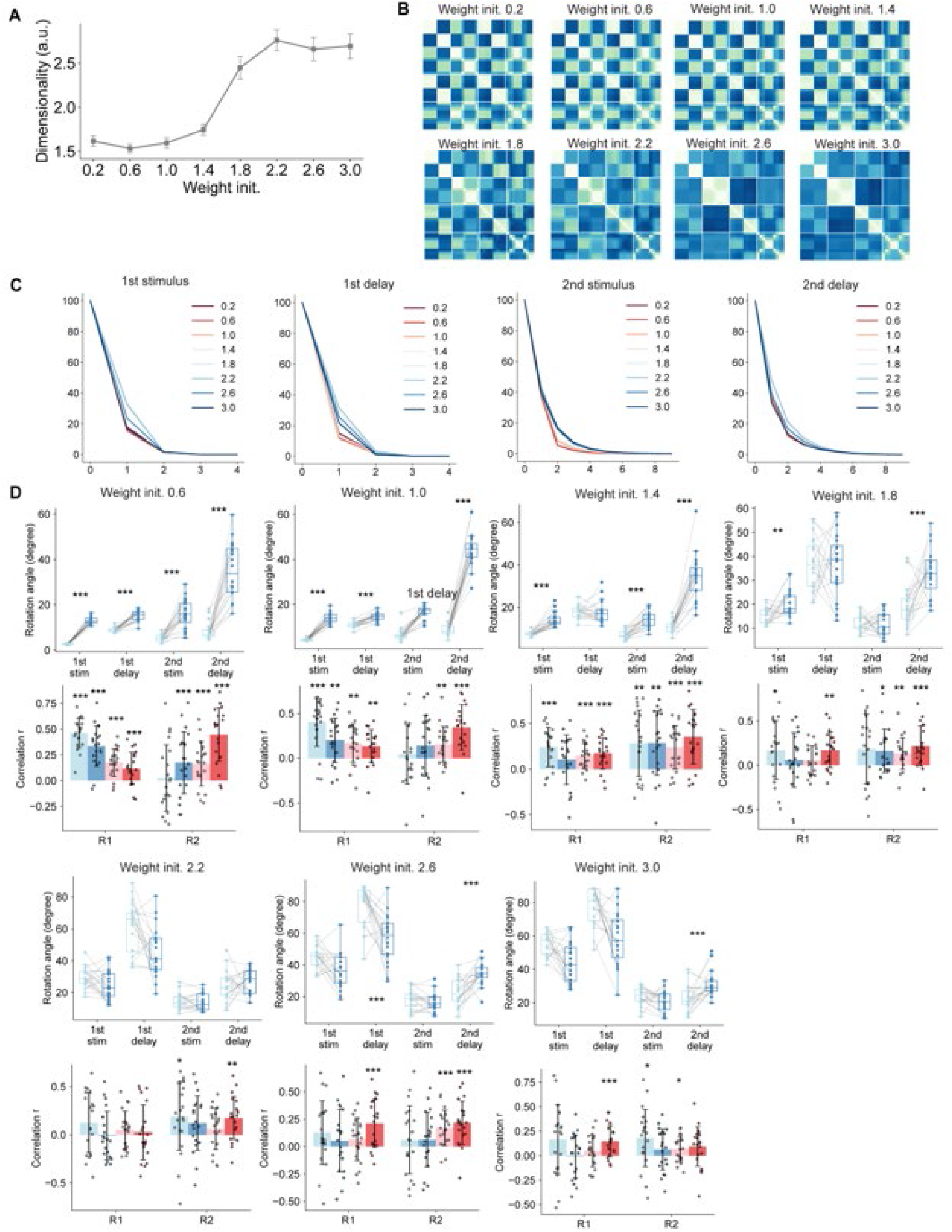
RNN modeling. (A) Neural activity dimensionality calculated by the participation ratio of the neural response during the delay period. (B) Representation dissimilarity matrices were calculated for recurrent neural networks (RNNs) with different training regimes. (C) Unexplained variance as a function of increasing principal components for each of the different task epochs for the different weight initialization. (D) The Beta-space subspace rotation angles of RNNs’ representations before and after the training, evaluated using a two-sided paired t-test (top) as well as the Pearson correlation *r* between the subspace rotation angles and the training performance (accuracy) of Remember 1^st^ (R1) and Remember 2^nd^ (R2) contexts, evaluated using a two-sided t-test (bottom). *: *p* < 0.05, **: *p* < 0.01, ***: *p* < 0.001.

